# Selenoprotein P Deficiency Drives Hepatocellular Carcinoma Progression via Induction of Neutrophil Senescence and Immunosuppressive Microenvironment

**DOI:** 10.1101/2025.06.24.661430

**Authors:** Jiazheng Jiao, Keying Xu, Kai Xiang, Rongrong Chen, Xiao Xu, Lixing Zhan

## Abstract

The tumor immune microenvironment (TIME) is critically shaped by immunosuppressive neutrophils, yet the molecular mechanisms underlying the functional reprogramming of these neutrophils remain poorly understood. Here, through integrative analysis of single-cell RNA sequencing (scRNA-seq) we showed the presence, in hepatocellular carcinoma (HCC) mice, of a novel subpopulation of senescent-like neutrophils that exhibit hepatic depletion of *Sepp1* in concert with the accumulation of the cellular senescence marker Cdkn1a, the immunosuppressive proteins S100a8 and 9, and the pro-angiogenic factor Vegfa. Furthermore, we demonstrated that the hepatic depletion of Sepp1, in combination with this unique subset of senescent neutrophils, drives immunosuppressive activity and further accelerates tumor growth. Mechanistically, the loss of tumor cell-derived Sepp1 impaired selenium uptake in tumor-infiltrating neutrophils (TINs) by disrupting Lrp8 receptor-mediated transport, thereby suppressing intracellular selenium metabolism. This metabolic perturbation led to reduced hydrogen selenide (H_2_Se) production and accumulation of S-adenosylmethionine (SAM), resulting in increased levels of trimethylation of lysine 4 on histone H3 protein (H3K4me3) modification in neutrophils and collectively establishing a pro-senescence chromatin landscape in tumor-associated neutrophils. Furthermore, we demonstrated that selenium supplementation not only restores Sepp1 expression but also is essential to reshaping anti-tumor immunity in HCC mice, including reversal of the senescent-like phenotype and providing synergistic effects with anti-Programmed cell death 1 (anti-PD-1) therapy. Collectively, our findings identify SEPP1 as a master regulator acting against neutrophil senescence and the potent immunosuppressive effects of senescence in HCC. We discuss the therapeutic potential of strategies to modulate senescent-like neutrophils and underscore the potential utility of selenium supplementation as an adjuvant to enhance immunotherapy efficacy in liver cancer.

## Introduction

The functional contributions of neutrophils in cancer are increasingly recognized, with both anti-tumor and pro-tumor roles reported^1–3^. The specific role of neutrophils in cancer is complex and controversial due to their high functional diversity and acute sensitivity to the microenvironment. This class of cells traditionally has been recognized as the first line of defense against infection and tissue injury, rapidly accumulating in large numbers at affected sites to fulfill protective roles^4^. Emerging evidence has revealed that neutrophils contribute to the generation of an immunosuppressive microenvironment, in which these cells act as key mediators of tumor initiation, tumor progression, tumor metastasis, and resistance to cancer therapies^5,6^. The interactions and coordination between these various neutrophil states and the cellular and molecular components of these cell, which govern the immunosuppressive microenvironment, remain incompletely understood.

The cellular senescence program is characterized by a permanent cell cycle arrest and a senescence-associated secretory phenotype (SASP). This growth-arrest mechanism operates primarily through transcriptional upregulation of the genes encoding cyclin-dependent kinase inhibitors, particularly cyclin-dependent kinase inhibitor 1A (CDKN1A; p21) and CDKN2A (p16)^7^. Indeed, expression of p21 is used as a marker of senescence^8,9^. There is growing recognition that the induction of neutrophil senescence by cell-intrinsic signals contributes to cancer initiation via the lingering presence of senescent cells in the tumor microenvironment (TME)^10^. A recent study reported that prostate tumor-derived apolipoprotein E (APOE) engages Triggering receptor expressed on myeloid cells 2 (TREM2) on neutrophils, inducing senescence and immunosuppressive phenotypes^11^. That study further demonstrated that depletion of senescent-like neutrophils provided marked suppression of the tumor burden.

Selenium is an essential element crucial for human health^12–15^. In biological systems, selenium is required for the biosynthesis of selenocysteine-containing selenoproteins that mediate its diverse, favorable effects on biology, including redox homeostasis, immunoregulation, and tumor suppression^16–19^.Among the 25 known human selenoproteins, selenoprotein P (SEPP1) is produced primarily by the liver and is the most abundant selenoprotein in plasma, accounting for approximately 50% of total serum selenium^16,17^. SEPP1 is a secreted protein composed of two distinct regions: a larger N-terminal domain containing a single selenocysteine residue within a redox motif, and a smaller C-terminal domain that harbors the remaining nine selenocysteines. SEPP1, as a carrier protein, plays an essential role in whole-body selenium homeostasis and distribution, distinct from the protein’s central role in selenium transport^20–24^. Current knowledge indicates that low-density lipoprotein receptor-related protein 8 (LRP8) is the established receptor for SEPP1, mediating its endocytosis to deliver selenium to target cells^25–29^. The binding of SEPP1 to LRP8 on target cells represents a critical step for supplying selenium to peripheral tissues. Numerous epidemiologic studies have demonstrated a significant correlation between the level of selenium in serum / toenail and the incidence of hepatocellular carcinoma (HCC)^30–33^.Furthermore, epidemiological data has demonstrated that it is higher serum SEPP1 levels are associated with a lower risk of HCC^34^. Despite these clinical observations, the mechanistic role of SEPP1 in hepatocarcinogenesis remains incompletely characterized. The immunomodulatory functions of selenium within the tumor microenvironment (TME)—particularly the role of SEPP1 in mediating tumor-immune crosstalk—remain poorly understood, despite their potential to shape immunosuppressive niches. this study investigated the role of selenium in liver cancer development, delineating a previously unrecognized role for selenium-SEPP1 transport in governing neutrophil senescence in HCC. We posited that SEPP1 deficiency creates a permissive niche for tumorigenesis through epigenetic reprogramming of neutrophils toward senescence-associated immunosuppression. Furthermore, our elucidation of the regulatory role of SEPP1 in senescent tumor-infiltrating neutrophils (TINs) unveiled novel immunomodulatory targets. These observations are expected to facilitate the development of combinatorial therapeutic strategies to enhance the clinical outcomes of immune therapies.

## Results

### Sepp1 is downregulated in various mice liver cancer contexts

To investigate the role of Sepp1 in HCC tumorigenesis, we employed a comprehensive approach using multiple murine models of primary liver cancer, including the classic model of HCC (via induction by diethylnitrosamine (DEN)) as well as six additional liver cancer models generated via hydrodynamic transfection with constructs encoding distinct combinations of oncoproteins, including NRas^G12V^, myristoylated protein kinase B (myr-AKT), c-Myc^T58A^, Yes-associated protein (Yap^S127A^), ΔN90β-catenin, and Notch intracellular domain 1 (NICD1) ^35,36^. Our data revealed that all seven HCC models exhibited pronounced depletion of the intracellular levels of Sepp1 compared to normal liver tissue (Figure S1a). Furthermore, serum Sepp1 levels were significantly decreased in mice transfected with constructs encoding NRas^G12V^ + myr-AKT, a widely utilized HCC model^3,37–40^, further corroborating the depletion of Sepp1 in HCC tissues (Figure S1b).

To this end, we established a series of mouse liver cancer cell lines with defined genetic alterations to identify the genes contributing to the decrease in Sepp1 levels. Primary hepatocytes were isolated from 6- to 8-week-old wild-type or *p53 tumor suppressor gene* (*Trp53*) -knockout mice using the classical two-step collagenase perfusion method to obtain highly active hepatocytes. These cells then were transfected with pT4 plasmids harboring oncogenes and SB100X plasmids. Since untransfected primary hepatocytes are unable to proliferate and undergo gradual cell death^41^, only transfected cells can proliferate and survive. Using this approach, we successfully established nine stable mouse liver cancer cell lines, each harboring a specific set of driver mutations (Figure S1c). Compared to primary mouse hepatocytes, all 9 of the newly established mouse liver cancer cells lines (PM:*Trp53*-null; c-Myc^T58A^ overexpressing, PR: *Trp53*-null; NRas^G12V^ overexpressing, PA: *Trp53*-null; myr-AKT overexpressing, PRA: *Trp53*-null; NRas^G12V^ and myr-AKT overexpressing, PMB: *Trp53*-null; c-Myc^T58A^ and ΔN90β-catenin overexpressing, PRB: *Trp53*-null; NRas^G12V^ and ΔN90β-catenin overexpressing, PMY: *Trp53*-null; c-Myc^T58A^ and Yap^S127A^ overexpressing, PAB: *Trp53*-null; myr-AKT and ΔN90β-catenin overexpressing, MR: c-Myc^T58A^ and NRas^G12V^ overexpressing) showed pronounced and consistent depletion of Sepp1, as was also seen in other tested HCC cell lines, including the previously described immortalized mouse hepatocyte cell lines AML12^42^, H22, and Hepa1-6 (Figure S1d). Collectively, these findings underscore the universal downregulation of Sepp1 in diverse murine liver cancer contexts, both in vivo and in vitro, highlighting the protein’s potential role as a critical molecular player in HCC pathogenesis.

### *Sepp1* deficiency exacerbates oncogene-driven liver cancer in mice

To investigate the role of Sepp1 in liver cancer progression, we employed a hydrodynamic tail vein injection (HDTVi) technique to induce liver tumors in mice. Liver tumors were obtained within 2 months after injection with constructs providing Sleeping Beauty transposon-mediated integration and *EF1a* promoter-driven expression of the oncoproteins NRas^G12V^ and myr-AKT. We further designed an expression knockdown strategy wherein two *Sepp1*-specific shRNAs were co-expressed simultaneously with NRas^G12V^ (Figure 1a). This approach enabled targeted depletion of *Sepp1* mRNA in tumor cells^37,43,44^. Quantitative PCR analysis using two independent primer sets confirmed that the levels of *Sepp1* mRNA were decreased by approximately 80% in the sh*Sepp1* group compared to the control group (Figure S2a). Western blot analysis further showed a substantial decrease in Sepp1 protein levels in the sh*Sepp1* group compared to the control group (Figure S2b).

**Figure 1:**
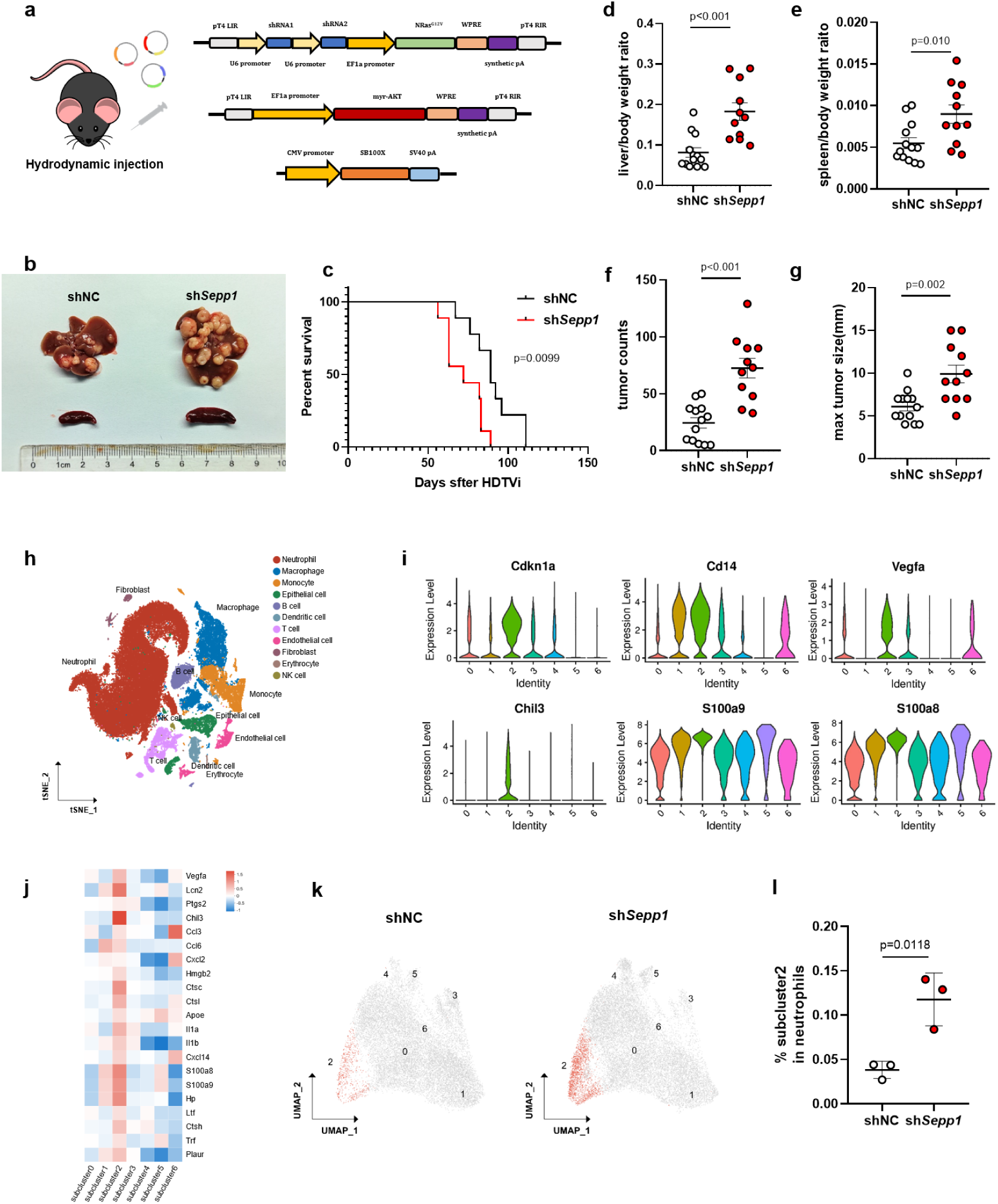
Sepp1 deficiency promotes oncogene-driven hepatocarcinogenesis and reveals a subset of senescent-like neutrophils. (a) Schematic representation of hydrodynamic tail vein injection (HDTVi) with constructs encoding oncoproteins (NRas^G12V^, myr-AKT) in combination with the *Sepp1* shRNA or with a control shRNA (shNC). (b) Representative necropsy images of livers from mice previously injected with shNC or sh*Sepp1*, demonstrating increased tumor burden in *Sepp1*-depeleted mice (compared to controls). (c) Kaplan-Meier survival curve showing significantly decreased survival in sh*Sepp1* mice compared to shNC controls (n=9 per group; p = 0.0099, as assessed using the log-rank test). (d, e) Quantification of liver-to-body weight ratio (d) and spleen-to-body weight ratio (e), showing a significant increase in both ratios for sh*Sepp1* mice compared to shNC mice (n = 11–13 per group). (f, g) Tumor counts (f) and maximum tumor size (g) in shNC versus sh*Sepp1* groups, showing a significantly higher tumor burden in the Sepp1-depleted group (n = 11–13 per group). (h) t-distributed stochastic neighbor embedding (t-SNE) plot of single-cell RNA sequencing data from liver tumors. (i) Violin plots showing expression levels of senescence- and immune suppression-related genes (*Cdkn1a*, *Cd14*, *Vegfa*, *Chil3*, *S100a9*, *S100a8*) across different cell types. (j) Heatmap showing the expression of selected factors, emphasizing that senescent-like Subcluster 2 exhibits high expression of secreted factors, including many SASP factors. (k) Uniform Manifold Approximation and Projection (UMAP) visualization of neutrophil clusters from shNC and sh*Sepp1* mice, showing that sh*Sepp1* samples are enriched for a distinct senescent-like neutrophile (Subcluster 2, marked in red). (l) Quantification of the percentage of cells in Subcluster 2 among neutrophils, showing a significant increase in the proportion of Subcluster-2 neutrophils in sh*Sepp1* mice compared to that in shNC controls (n = 3 mice per group). Data are presented as mean ± SEM; where not specified, statistical significance was determined using a two-tailed unpaired Student’s t-test.

Kaplan-Meier survival analysis revealed a significant decrease in overall survival in the sh*Sepp1* group compared to the negative control (shNC) group (Figure 2c). Additionally, the liver-to-body weight ratio and spleen-to-body weight ratio were significantly increased in the sh*Sepp1* group compared to controls (Figures 1b, d, e). Tumor quantification demonstrated that *Sepp1* knockdown resulted in an increase in both the number of tumors and the maximum tumor size (Figures 1f, g). Additionally, serum levels of aspartate aminotransferase (AST) and alanine aminotransferase (ALT), markers of liver damage, were significantly elevated in the sh*Sepp1* group compared to the control group (Figures S2c, d). Furthermore, serum glutathione peroxidase (GPx) activity, a surrogate marker of systemic selenium status, was significantly decreased in the sh*Sepp1* group compared to the control group (Figure S1e).

**Figure 2:**
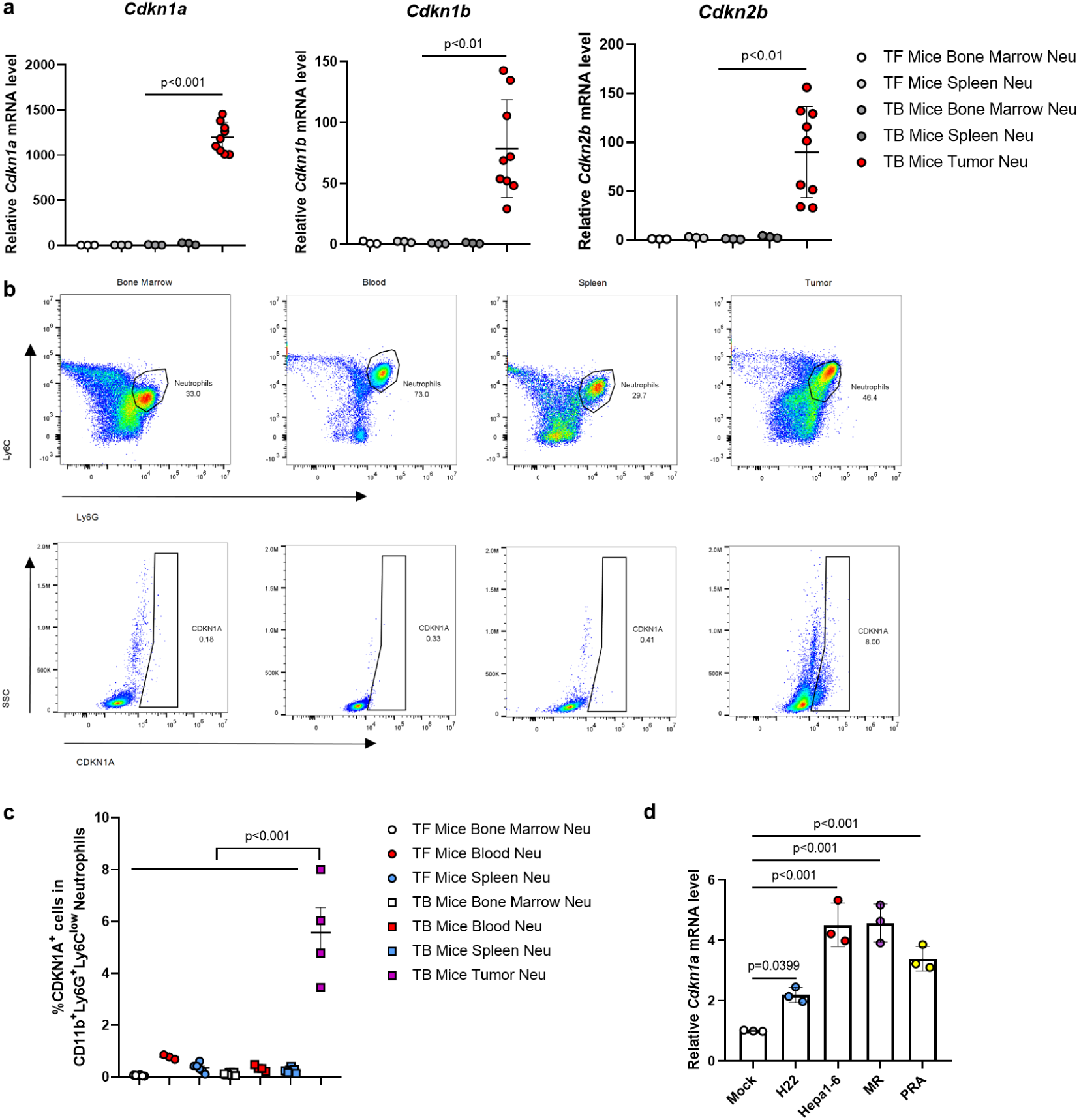
p21^+^ senescent-like neutrophils are present exclusively in tumor tissues. (a) Quantitative RT-PCR analysis of gene expression in neutrophils isolated from various tissues from tumor-free (TF) and tumor-bearing (TB) mice, including bone marrow (BM), spleen, and tumor. Markers of senescence, including *Cdkn1a*, *Cdkn1b*, and *Cdkn2b*, were selectively and significantly elevated in tumor-associated neutrophils, indicating a distinct senescent-like phenotype within the tumor microenvironment (n = 3–9 per group). (b) Flow cytometric analysis showing the proportion of neutrophils from different tissue sources, including BM, blood, spleen, and tumors, in TB mice, showing that p21-positive neutrophils are exclusively present in tumor-infiltrating neutrophils (TINs). (c) Quantification of the percentage of p21-positive neutrophils from different tissue sources, including BM, blood, spleen, and tumor, in TF and TB mice. TINs exhibit a significantly higher percentage of p21-positive cells compared to neutrophils from other tissue sources and TF conditions (n = 3–8 per group). (d) Quantitative RT-PCR analysis of gene expression in primary neutrophils treated with conditioned medium from liver cancer cell lines. The level of *Cdkn1a* mRNA was significantly elevated in neutrophils treated with H22, MR, PRA, and Hepa1-6 conditioned medium compared to the mock control (n = 3 per group). Data are presented as mean ± SEM; statistical significance was determined using One-way ANOVA with post hoc Dunnett’s multiple comparisons tests.

We extended this analysis by conducting rescue experiments in which we constructed and expressed a *Sepp1* rescue plasmid in the sh*Sepp1* background (Figure S2f, g). Restoration of *Sepp1* expression successfully reversed the severe tumor phenotypes, including decreasing liver-to-body weight ratios and decreasing tumor counts compared to controls (Figures S2h-j). Together, these results demonstrated that Sepp1 plays a protective role in liver cancer.

### scRNA-seq identifies a distinct subpopulation of senescent-like neutrophils implicated in HCC

To elucidate the mechanistic role of Sepp1 in HCC progression and the functional impact of the protein within the TME, we performed single-cell RNA sequencing (scRNA-seq) on tumor samples derived from *Sepp1* knockdown (n=3) and shNC control (n=3) mice. Additionally, two samples each of tumors derived from mice that lacked the shRNA expression cassette, mice that had been treated with selenium supplementation, and corresponding controls, were included in the analysis. A total of 90031 cells from 8 samples were comprehensively profiled.

The t-distributed stochastic neighbor embedding (t-SNE) plot (Figures 1h, S3a) revealed distinct cellular populations within the TME, with neutrophils forming a prominent cluster. Subsequent reclustering of neutrophils identified seven distinct sub-populations (Figure S3b), among which Subcluster 2 exhibited a unique transcriptional signature. Violin plots of gene expression (Figure 1i) revealed that Subcluster 2 was characterized by elevated levels of the *Cdkn1a* transcript (a marker of cellular senescence), along with increased levels of *S100a8*, *S100a9*, and *Vascular endothelial growth factor A* (*Vegfa)* mRNAs (encoding proteins associated with tumor progression, immunosuppression, and angiogenesis). Furthermore, altered expression patterns of *Chitinase-like 3* (*Chil3*) and *Cluster of differentiation 14* (*Cd14*) transcripts in cells of this subcluster suggested a functional reprogramming of neutrophils towards an immunosuppressive phenotype, consistent with the results of prior studies^6,45^.

Analysis, across the seven neutrophil subclusters, of the expression of genes encoding secreted proteins highlighted a robust SASP signature in Subcluster 2. Transcripts encoding classical SASP factors^46,47^, including *Interleukin 1a* (*Il1a)*, *Il1b*, *Chemokine (C-C motif) ligand 3* (*Ccl3*), *Ccl6*, *Vegfa*, *Prostaglandin-endoperoxide synthase 2* (*Ptgs2*), *S100a8*, and *S100a9*, were significantly upregulated in Subcluster 2 compared to other subclusters (Figure 1j). Notably, *Plasminogen activator urokinase surface receptor* (*Plaur*), which encodes a cell-surface protein shown to be broadly induced during senescence^48,49^, also accumulated to significantly higher levels in Subcluster 2 than in other subclusters. Considered together, these results identify Subcluster 2 as a group of senescent-like neutrophils. Intriguingly *Sepp1* knockdown resulted in a significant expansion of Subcluster 2 within the neutrophil compartment compared to that in shNC mice (Figures 1k, l), suggesting a potential role for *Sepp1* in regulating neutrophil senescence. Gene Ontology (GO) enrichment analysis of upregulated signature genes in Subcluster 2 revealed enrichment in loci associated with neutrophil chemotaxis and inflammatory responses, underscoring the functional relevance of this subset in HCC progression (Figure S3c).

We employed Monocle3, an analysis toolkit for single-cell RNA-seq data, to perform pseudotime analysis on all neutrophils. This test revealed that Subcluster 2 occupies the terminal stage of the pseudotime trajectory, indicating that this class of neutrophils represents a distinct differentiation state compared to other subclusters (Figure S3d). This finding aligns with the senescent-like characteristics observed for Subcluster 2, as evidenced by the expression of senescence-associated markers. Gene expression dynamics along the pseudotime trajectory (Figures S3e-f) provided critical insights into the molecular programs governing these transitions. Notably, *Cdkn1a* exhibited a pronounced upregulation in the later stages of pseudotime, particularly in Subcluster 2, further corroborating the identification of this subcluster as a senescent population. Additionally, the expression of *Vegfa* and *Chil3* also increased progressively along the trajectory; in contrast, the expression of *Cd14* and *S100a8/9* displayed oscillatory patterns with peak expression at the terminal stages. These patterns of expression are consistent with the role of these genes in immune modulation and tumor promoting function.

To functionally validate the role of neutrophils in *Sepp1*-knockdown liver tumors, we performed neutrophil depletion experiments in an independent experiment. Neutrophil depletion provided decreased tumor burden in *Sepp1*-knockdown liver tumor samples (Figures S4a-d). Furthermore, flow cytometric analysis revealed that neutrophil depletion leads to a significant increase in the proportion of T cells and natural killer (NK) cells within the CD45^+^ immune cell population (Figures S4e, f). These findings suggested that neutrophil depletion alleviates the immunosuppressive state within the TME, thereby restoring anti-tumor immunity and relieving tumor progression. This inference is consistent with the results of previous research^50^ that highlighted the critical role of neutrophils in shaping immunosuppressive microenvironments in cancer.

Thus, our results demonstrated that Sepp1 deficiency in tumor cells drives the expansion of a senescent-like neutrophil subpopulation characterized by elevated expression of senescence markers and pro-tumorigenic factors. We hypothesize that this subset of neutrophils contributes to tumor progression by fostering an immunosuppressive microenvironment and promoting angiogenesis. These findings provide novel insights into the interplay between Sepp1, neutrophil senescence, and tumor immunity, offering potential therapeutic avenues for targeting neutrophil-mediated immunosuppression in HCC.

### p21-positive neutrophils are found predominantly in tumor tissues

Neutrophils, which are characterized by a short lifespan and exhibit remarkable heterogeneity and plasticity, are known to play diverse roles in pathological contexts. To investigate whether senescent-like neutrophils are specifically enriched within tumors, we isolated neutrophils from the bone marrow (BM), spleen, and tumor tissues of healthy mice and HCC-bearing animals using magnetic-activated cell sorting (MACS). Transcriptomic analysis of these tissues revealed that TINs exhibited significantly elevated expression of classic immunosuppressive genes, including Arginase 1 (*Arg1*), *Nitric oxide synthase 2* (*Nos2*), *Cd14*, *Ptgs2*, and *Cd274* (compared to neutrophils from non-tumor tissues) (Figure S5a), suggesting that these cells undergo functional reprogramming within the TME. Notably, TINs also displayed significant upregulation of senescence-associated markers, including *Cdkn1a*, *Cdkn1b*, and *Cdkn2b*, compared to neutrophils isolated from the BM or spleen of either healthy or tumor-bearing mice (Figure 2a). Moreover, *Cdkn2a* transcripts also accumulated to significantly higher levels in TINs compared to BM-derived neutrophils, although the magnitude of the increased expression of *Cdkn2a* was much lower than that observed for *Cdkn1a* in the same cells (Figure S5a).

Next, we utilized flow cytometry to assess the proportion of p21-positive neutrophils in the BM, spleen, blood, and tumor tissues of healthy and HCC-bearing mice. Consistent with the transcriptional data, p21^+^ neutrophils were detected predominantly within tumor tissues, with minimal presence in other compartments (Figures 2b, c).

To verify whether this phenomenon also occurs in human samples, we examined single-cell sequencing data obtained for neutrophils derived from human liver cancer samples^43^ (http://meta-cancer.cn:3838/scPLC). Notably, analysis of neutrophil subsets within this data set revealed that the expression of *CDKN1A* (encoding the human p21 homolog) also was found predominantly in human TINs, whereas expression of this gene was much lower in human neutrophils from peripheral blood or peritumoral tissues (Figure S5b).

The selective accumulation of senescent neutrophils within tumors raises the question of whether the TME drives this phenomenon. To address this issue, we incubated mouse neutrophils with the supernatants from various mouse liver cancer cell lines. Using Xite™ Red beta-D-galactopyranoside to detect senescence-associated beta-galactosidase (SA-β-GAL; a marker of senescence^51^), we demonstrated that exposure to the supernatants from multiple liver cancer cell lines increased the proportion of SA-β-GAL^+^ neutrophils compared to the effect of exposure to control medium (Figures S5c, d). We also found that the cell supernatants induced p21 expression, along with the accumulation of *Cd14*, *Vegfa*, and *Chil3* transcripts, all of which correspond to the profile of neutrophils of Subcluster 2 (Figures 2d, S4e-g). Taken together, these data indicated that in both mouse and human, tumor-associated neutrophils (TANs) exhibit a distinct senescence phenotype characterized by elevated p21 expression, a feature not observed in neutrophils from BM, blood, or spleen.

### Sepp1 deficiency induces neutrophil senescence and immunosuppression, leading to reduced T cell infiltration and activity in tumors

Next, we conducted comprehensive analyses of immune cell populations and gene expression profiles in sh*Sepp1* versus shNC mouse liver tumor models. Flow cytometric analysis identified a significant increase in the percentage of neutrophils in Sepp1-depeleted tumors compared to control tumors (Figure 3a, S6a). TINs were isolated using MACS, and quantitative reverse transcriptase-polymerase chain reaction (RT-qPCR) analysis revealed significantly increased accumulation, in the *Sepp1*-knockdown group (compared to controls), of transcripts encoding senescence markers, including such as *Cdkn1a*, *Cdkn2a*, *Cdkn2b*, and *Cdkn2d* (Figures 3b, S6b, c). Additionally, we observed significantly elevated levels (in the *Sepp1*-knockdown group compared to controls) of transcripts encoding immunosuppressive markers (including *S100a8/9*, *Cd14*, and *Chil3*) and pro-angiogenic markers (*Vegfa*) (Figures 3c, d). These results corroborated the findings obtained via scRNA-seq. Additionally, flow cytometric analysis demonstrated significantly increased protein levels of S100a8/9 in neutrophils from the *Sepp1*-knockdown group compared to those from controls (Figure 3e).

**Figure 3:**
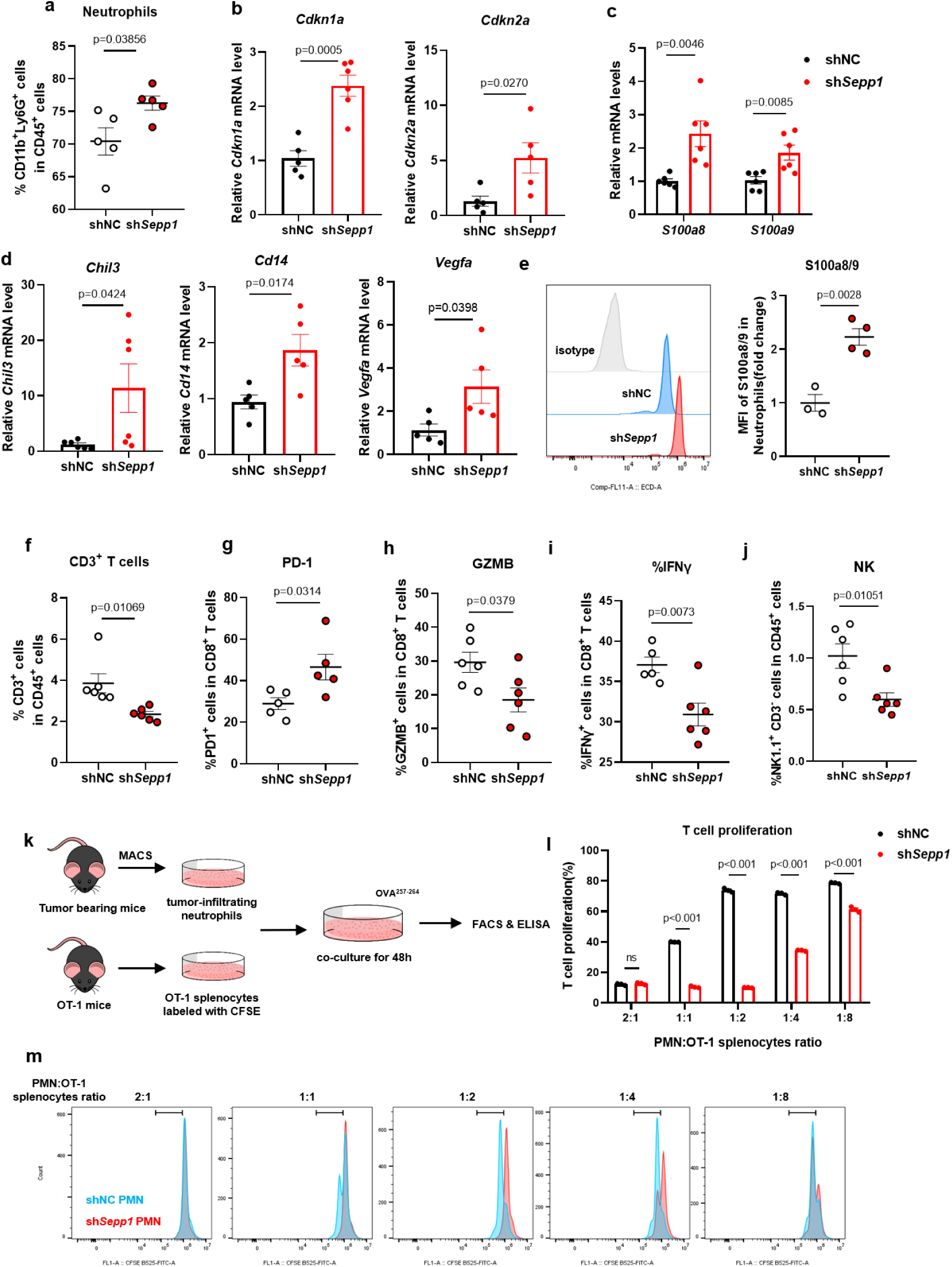
Sepp1 depletion induces neutrophil senescence and immunosuppression, reducing T cell infiltration and activity in tumors. (a) Flow cytometric analysis showing an increased percentage of neutrophils (CD11b^+^ lymphocyte antigen 6 family member G-positive (Ly6G^+^) cells) in the tumor microenvironment of sh*Sepp1* mice compared to shNC controls (n = 5–6 per group). (b) Quantitative RT-PCR analysis of senescence-associated markers *Cdkn1a* and *Cdkn2a* in tumor-infiltrating neutrophils (TINs) (n = 5–6 per group). (c) mRNA expression levels of *S100a8* and *S100a9* were significantly elevated in TINs from sh*Sepp1* mice compared to shNC controls (n = 6 per group). (d) mRNA expression levels of *Chil3*, *Cd14*, and *Vegfa* were significantly elevated in TINs from sh*Sepp1* mice compared to shNC controls (n = 5-6 per group). (e) Flow cytometric analysis of S100A8/9 protein expression in neutrophils, with representative histogram overlay (left) and quantification of mean fluorescence intensity (MFI) (right) (n = 3 per group). (f) Flow cytometric quantification of CD3^+^ T cells in the tumor microenvironment of shNC and sh*Sepp1* mice, indicating significantly decreased T cell infiltration in sh*Sepp1* tumors compared to controls (n = 6 per group). (g–i) Functional analysis of CD8^+^ T cells in tumors, demonstrating increased programmed cell death 1 (PD-1) expression (g), decreased granzyme B (GZMB) levels (h), and decreased interferon gamma (IFNγ) levels (i) in sh*Sepp1* mice compared to controls (n = 5–6 per group). (j) The percentage of natural killer (NK) cells (NK1.1^+^ CD3^−^) was significantly decreased in the tumor microenvironment of sh*Sepp1* mice compared to shNC animals (n = 6 per group). (k) Schematic representation of the experimental setup for assessing T cell suppression by TINs. Neutrophils were isolated from the tumors of mice injected with shNC or sh*Sepp1* constructs using Magnetic Activated Cell Sorting (MACS). These neutrophils were co-cultured with CFSE-labeled OT-1 splenocytes from OT-1 mice at various ratios (2:1 to 1:8) in the presence of OVA_257−264_ peptide for 48 hours, followed by analysis using FACS and ELISA. (l) T cell proliferation analysis measuring the percentage of proliferating OT-1 T cells co-cultured with neutrophils isolated from shNC or sh*Sepp1* tumors. This analysis showed significantly decreased T cell proliferation in the presence of neutrophils from *Sepp1*-depleted tumors across different neutrophil: OT-1 ratios (compared to those cultured in the presence of neutrophils from shNC tumors) (n = 3 per group). (m) Representative flow cytometry histograms of CFSE dilution in OT-1 splenocytes, illustrating T cell proliferation at different neutrophil: OT-1 splenocyte ratios. Data are presented as mean ± SEM; statistical significance was determined using two-tailed unpaired Student’s t-tests.

In parallel, we observed a significant decrease in the proportion of total CD3^+^ T cells in the sh*Sepp1* group (Figure 3f, S6d) compared to control animals. Both CD4^+^ and CD8^+^ T cell populations were significantly decreased in *Sepp1*-depeleted tumors compared to control tumors (Figures S6e, f). To further assess the functional status of CD8^+^ T cells, we analyzed their exhaustion and activation markers. *Sepp1* knockdown resulted in a significant increase in the percentage of Programmed cell death 1-positive (PD-1^+^) CD8^+^ T cells (Figures 3g, S6h). Concurrently, we observed decreased expression of granzyme B (GZMB) and interferon gamma (IFNγ) (Figures 3h, i, S6h), indicating decreased cytotoxic activity of CD8^+^ T cells in Sepp1-depeleted tumors compared to control tumors. Furthermore, the proportion of NK cells was significantly diminished in the sh*Sepp1* group compared to the control (Figure 3j), suggesting a broad suppression of cytotoxic immune responses in the absence of Sepp1.

To elucidate the functional impact of neutrophil senescence on T cell-mediated immunoactivity, we performed co-culture experiments using TINs isolated from Sepp1-depeleted and control tumors. TINs were purified via MACS and co-cultured with carboxyfluorescein succinimidyl ester (CFSE) -labeled OT-1 splenocytes in the presence of an ovalbumin-derived peptide (OVA_257-264_) for 48 hours to assess T cell proliferation and cytokine secretion profiles (Figure 3k).

Flow cytometric analysis revealed a significant impairment in the proliferative capacity of OT-1 CD8^+^ T cells when co-cultured with TINs from sh*Sepp1* tumors compared to controls (Figures 3l, m). This observation implied that TINs from *Sepp1* knockdown tumors possess enhanced immunosuppressive properties, presumably via impaired T cell activation and proliferation. Further assessment of cytokine production showed a significant reduction in the levels of Ifnγ and Il2 in the presence of TINs from *Sepp1* knockdown tumors (Figures S6i, j).

Using another HCC mouse models constructed by double-knockout (DKO) of *Trp53* and *Phosphatase and tensin homolog* (*Pten*) (Figures S7a, b), we again observed that the further knockout of *Sepp1* (the triple-knockout; TKO) significantly increased the hepatic tumor burden compared to that in DKO (Figure S7c). Tumor quantification demonstrated that *Sepp1* knockout resulted in increases in the number of tumors, the liver-to-body weight ratio, and the spleen-to-body weight ratio (Figures S7d-f). Flow cytometric analysis also revealed an increase in the proportion of senescent neutrophils, as indicated by higher SA-β-GAL mean fluorescence intensity (MFI) (Figure S7g). T cell analysis showed a decrease in the proportion of total CD3^+^ T cells in the TKO group compared to the DKO parent (Figure S7h). Both CD4^+^ and CD8^+^ T cell populations also were significantly decreased in TKO compared to DKO (Figures S7i, j). Additionally, the frequency of PD-1⁺ exhausted CD8⁺ T cells was significantly increased and that of IFN-γ⁺ CD8⁺ T cells was significantly decreased, in TKO compared to DKO (Figures S7k, l). Next, TINs were isolated using MACS and used to isolate RNA; RT-qPCR analysis revealed significantly increased levels of senescence marker mRNAs such as *Cdkn1a* and *Cdkn2d* in the TKO group compared to the controls, Additionally, we observed elevated levels of immunosuppressive marker transcripts like *S100a8/9*, *Cd14,* and *Chil3*, corroborating our findings in the NRas^G12V^ + myr-AKT mouse model (Figures S7m-r).

Overall, Sepp1 deficiency led to an increase in the proportion of senescent-like neutrophils and promoted T cell exhaustion and attenuation of cytotoxic CD8^+^ T cell activity. These results underscore the critical role of Sepp1 in modulating neutrophil-T cell crosstalk and highlight this protein’s potential as a therapeutic target to restore antitumor immunity.

### Selenium exerts antitumor effects in a Sepp1-dependent manner

To verify whether selenium supplementation promotes antitumor activity in a Sepp1-dependent manner, we treated tumor-bearing *Sepp1* knockdown mice and control mice by providing the animals with ad libitum access to drinking water containing 10 ppm of sodium selenite or selenomethionine. While both organic and inorganic selenium supplementation significantly inhibited tumor progression, this effect was abrogated in *Sepp1* knockdown models compared to the control mice (Figures 4a-e, S8b). Enzyme-linked immunosorbent analysis (ELISA) confirmed that, when provided with selenium supplementation, the controls (but not the sh*Sepp1* mice) exhibited elevated serum levels of Sepp1 (Figure 4f), consistent with changes in the tumors (Figure S8a). Supplementation with selenium attenuated the levels of *Cdkn1a* and *S100a9* transcripts in neutrophils (Figures 4h, i). However, in tumors with *Sepp1* knockdown, selenium supplementation did not have this effect. Assessment of the serum level of S100a9 protein provided results that were consistent with the RT-qPCR data (Figure 4g).

**Figure 4:**
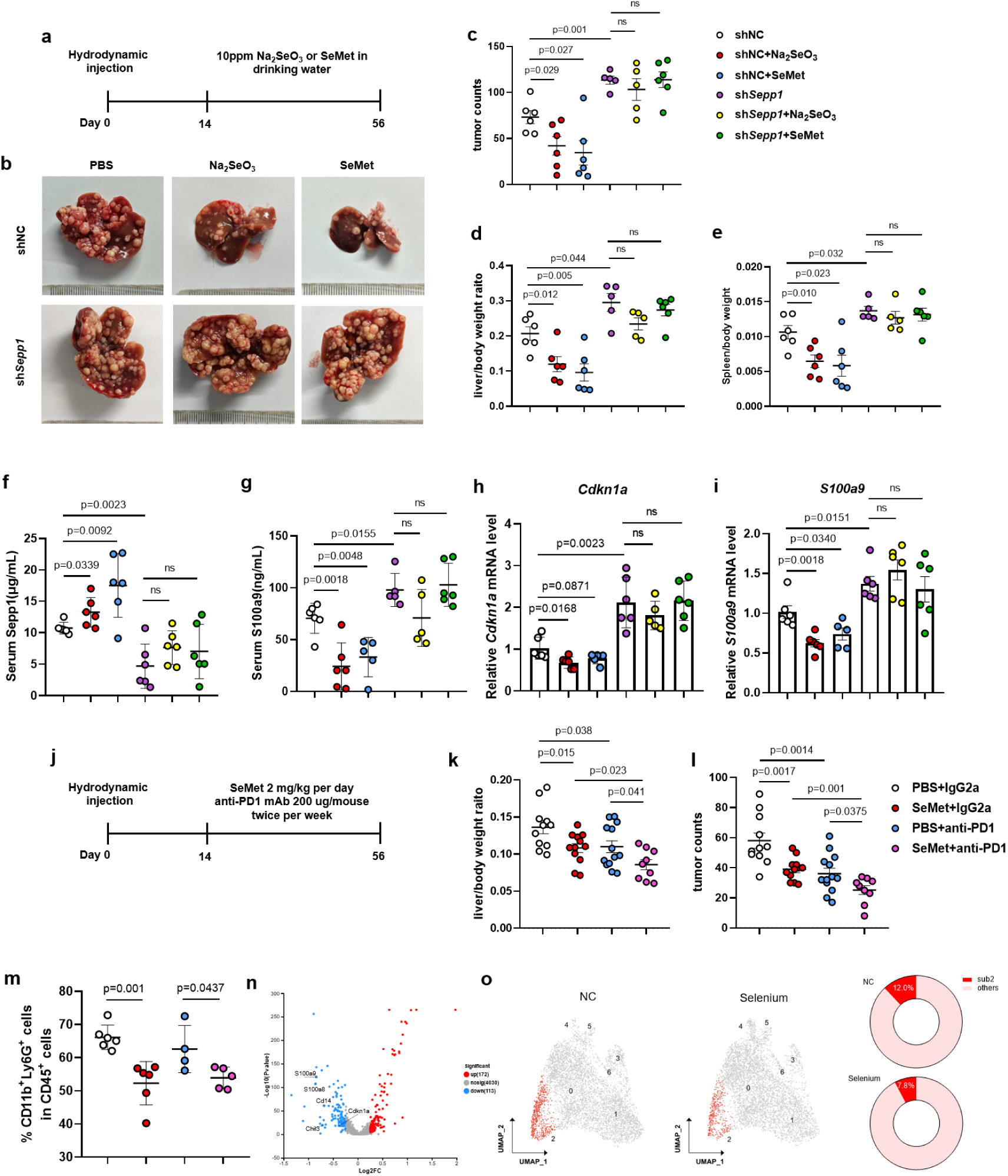
Selenium inhibits tumor growth and exhibits synergistic effects with anti-PD-1 in a Sepp1-dependent manner. (a) Experimental timeline for selenium treatment. Mice were injected via hydrodynamic tail vein (HDTVi) with constructs encoding oncogenic NRas^G12V^ and myr-AKT proteins, along with shNC or sh*Sepp1* constructs, and provided (from Day 14 to Day 56 post HDTVi) with ad libitum access to drinking water containing 10 ppm sodium selenite (Na_2_SeO_3_) or selenomethionine (SeMet). (b) Representative images of livers from selected groups. sh*Sepp1* mice exhibited more extensive tumor growth than did shNC mice. Selenium treatment reduced the tumor burden in shNC mice, but had little effect on the severe tumor burden in sh*Sepp1* mice. (c – e) Tumor count (c), liver-to-body weight ratio (d), and spleen-to-body weight ratio (e) measurements. sh*Sepp1* tumors exhibited significantly higher tumor burden than did shNC controls, as indicated by increased tumor counts, relative liver weight, and relative spleen weight. Selenium treatment significantly attenuated the increases in tumor counts and weight ratios in shNC mice, but had no significant effect on the severe tumor burden observed in sh*Sepp1* mice (n = 5-6 per group, p-values as indicated). (f, g) Serum Sepp1 (f) and S100A9 (g) levels measured by ELISA, showing significantly increased Sepp1 levels and decreased S100A9 levels in selenium-treated shNC mice, whereas selenium treatment did not significantly alter these markers in sh*Sepp1* mice (n = 6 per group, p-values as indicated). (h, i) Quantitative RT-PCR analysis of *Cdkn1a* (h) and *S100a9* (i) in tumor-infiltrating neutrophils (TINs) from treated mice. TINs from sh*Sepp1* tumors exhibited elevated levels of these markers compared to those from shNC animals. Selenium treatment was associated with decreased levels of *Cdkn1a* and S100a9 transcripts in neutrophils from shNC tumors, but had no significant effect on the levels of these marker in neutrophils from sh*Sepp1* tumors (n = 5–6 per group). (j) Timeline schematic for combination treatment with SeMet and anti-PD-1 antibody. Mice received HDTVi with constructs encoding oncogenic NRas^G12V^ and myr-AKT proteins, and then were treated (from Day 14 to Day 56 post HDTVi) with SeMet (2 mg/kg per day) and/or anti-PD-1 monoclonal antibody (200 μg/mouse, twice per week). (k, l) Liver-to-body weight ratios (k) and tumor counts (l), demonstrating significant reduction with combination therapy of SeMet and anti-PD-1 compared to monotherapy or control treatments and indicating the synergistic effect of selenium supplementation with immune checkpoint blockade (n = 9–13 per group, p-values as indicated). (m) Flow cytometric analysis showing the percentage of neutrophils in tumor tissues from treatment groups, demonstrating a decrease in neutrophil proportion with SeMet therapy (n = 4–6 per group). (n) Volcano plot of differentially expressed genes in neutrophils from SeMet-treated versus control mice, highlighting that selenium supplementation is associated with decreased transcript levels for key genes including *Cdkn1a*, *S100a8/9*, *Chil3*, and *Cd14*. (o) Uniform Manifold Approximation and Projection (UMAP) plots of TINs from phosphate-buffered saline (PBS; vehicle) and SeMet-treated mice, showing that the distinct senescent-like Subcluster 2 (marked in red) decreased after selenium supplementation. Data are presented as mean ± SEM; statistical significance was determined using two-tailed unpaired Student’s t-test.

As demonstrated above, tumors with *Sepp1* knockdown showed a higher proportion of aging subgroups and exhibited stronger immunosuppressive abilities in vitro (compared to control tumors). Since immunosuppressive neutrophils are the main negative regulators of antitumor immunity and are considered a primary reason for poor efficacy of, or resistance to, immunotherapy^52–54^, this observation raised the questions of (a) whether selenium enhances the efficacy of anti-PD-1 immune checkpoint inhibitors when used in combination, and (b) whether this effect is mediated through Sepp1. To evaluate whether selenium supplementation improved the efficacy of immunotherapy, we combined selenomethionine exposure with anti-PD-1 therapy in liver cancer mice (Figure 4j). Notably, the combination of selenium supplementation and anti-PD-1 antibody resulted in a significant attenuation of the tumor burden compared to control (untreated) mice or those receiving the individual treatments (Figures 4k, l, S8c). These results suggested a synergistic effect between selenium supplementation and immune checkpoint blockade. Flow cytometric analysis of immune cell populations in these animals demonstrated a significant decrease in the proportion of neutrophils in the selenium supplementation + antibody group compared to control (untreated) mice or those receiving the individual treatments (Figure 4m). Single-cell RNA sequencing revealed selenium-associated attenuation of the transcript levels of key senescent-like neutrophil genes (*Cdkn1a*, *S100a8/9*, *Chil3*, and *Cd14*) compared to control neutrophils (Figure 4n). The t-SNE plots revealed that selenium supplementation led to attenuation of the expansion of the senescent-like neutrophil Subcluster 2 (Figure 4o). These findings confirm that Sepp1 serves as a crucial mediator of selenium’s ability to enhance immunotherapy by modulating neutrophil function and alleviating tumor-induced immunosuppression

### Targeting senescent neutrophils reverses immunosuppression in Sepp1-deficient liver cancer

Here, we employed a well-established senolytic regimen combining dasatinib (5 mg/kg) and quercetin (50 mg/kg) ^55,56^ (D& Q) to test whether the classic senolytic treatment mitigated the effects of Sepp1 deficiency. Treatment was initiated 14 days post-hydrodynamic injection (tumor induction) and administered twice weekly for six weeks (Figure 5a). We observed that D&Q treatment significantly decreased the tumor burden in sh*Sepp1* mice compared to control mice (Figures 5b, c, S9a, b).

**Figure 5:**
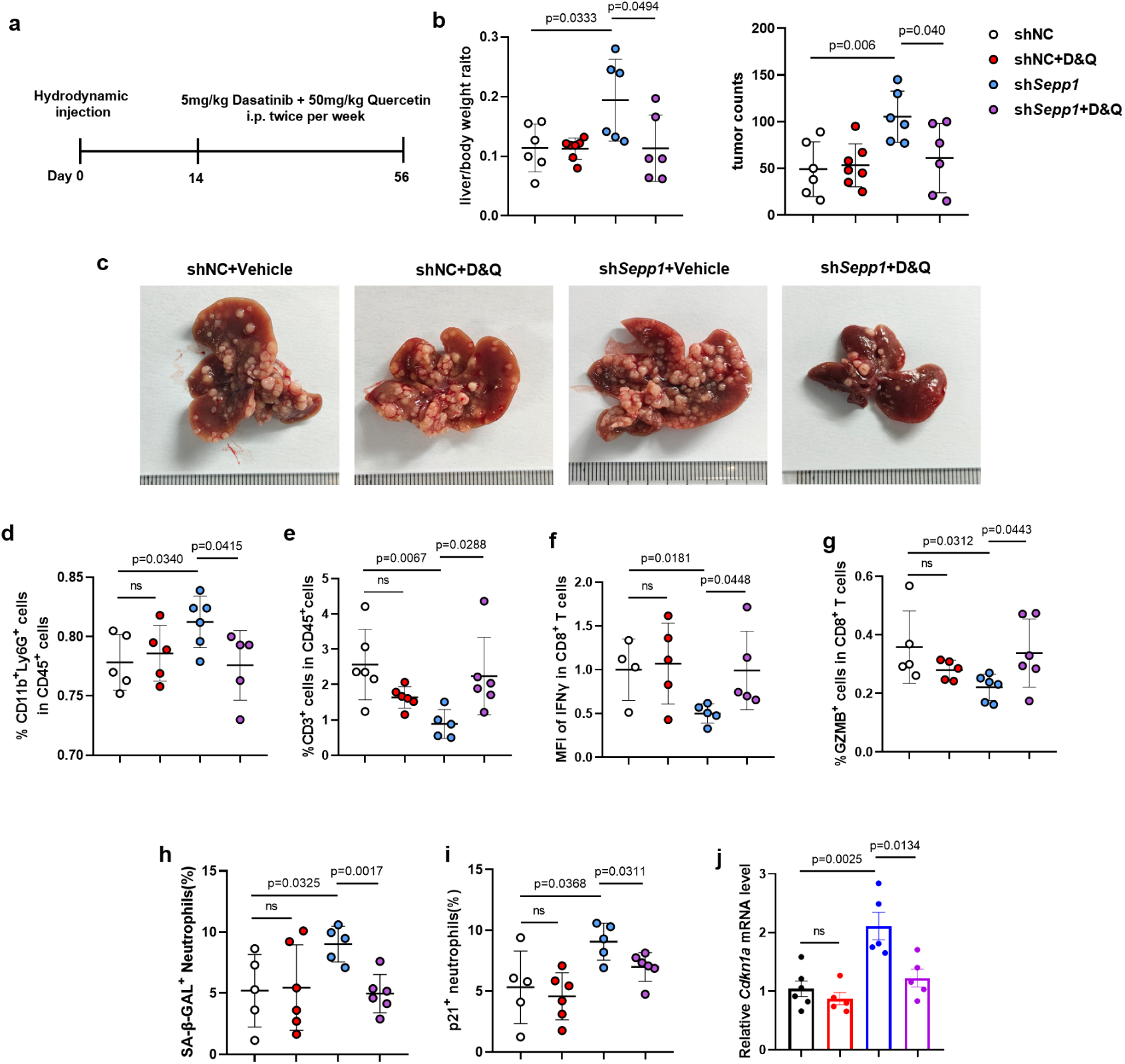
Targeting neutrophil senescence to mitigate Sepp1 deficiency in liver cancer. (a) Experimental timeline for dasatinib and quercetin (D&Q) treatment. Mice were injected via hydrodynamic tail vein (HDTVi) with constructs encoding oncogenic NRas^G12V^ and myr-AKT proteins along with shNC or sh*Sepp1* constructs, and subsequently treated (from Day 14 to Day 56 post HDTVi) with the combination of D&Q (5 and 50 mg/kg, respectively) by twice-weekly intraperitoneal injection. (b-c) Liver-to-body weight ratio and tumor counts indicating that sh*Sepp1* mice had increased tumor burden compared to shNC. Treatment with D&Q significantly attenuated liver-to-body weight ratios and tumor counts in sh*Sepp1* mice (n = 6–7 per group). (d–g) Flow cytometric analysis showing the percentage of CD11b^+^ lymphocyte antigen 6 family member G-positive (Ly6G^+^) neutrophils (d), CD8+ T cells (e), Mean fluorescence intensity (MFI) of IFNγ in CD8^+^ T cells (f), and percentage of granzyme B-positive (GZMB^+^) CD8^+^ T cells (g) in tumor tissues. D&Q treatment significantly attenuated neutrophil infiltration (d) and potentiated CD8^+^ T cell activation markers (e–g) in sh*Sepp1* tumors (n = 4–6 per group). (h, i) Flow cytometric analysis of senescence markers in tumor-infiltrating neutrophils (TINs) from treated mice. The proportions of senescence-associated β-galactosidase (SA-β-Gal) -positive neutrophils (h) and p21^+^ neutrophils (i) were significantly reduced in D&Q-treated sh*Sepp1* mice, indicating effective senescence targeting (n = 5-6 per group). (j) Quantitative RT-PCR analysis of *Cdkn1a* in TINs, demonstrating reduced expression of senescence-associated *Cdkn1a* in D&Q-treated sh*Sepp1* mice (n = 5–6 per group). Data are presented as mean ± SEM; statistical significance was determined using two-tailed unpaired Student’s t-tests.

Immune profiling revealed that D&Q treatment not only decreased the proportion of TINs in Sepp1-depeleted mice (Figure 5d), but also significantly enhanced T cell-mediated antitumor immunity, as indicated by the increased MFI of IFNγ and an elevated percentage of GZMB^+^ CD8^+^ T cells in the sh*Sepp1* group compared to the control animals (Figures 5e–g). Flow cytometric analysis showed a significant attenuation in the proportion of SA-β-Gal positive neutrophils and of p21^+^ neutrophils in the D&Q-treated sh*Sepp1* group (compared to untreated mice) (Figures 5h, i). These findings were further corroborated by RT-qPCR, which showed attenuation of senescence-associated markers (*Cdkn1a* and *Cdkn2a* transcripts) in TINs following D&Q treatment (Figures 5j, S9c). Additionally, D&Q treatment was associated with decreases (compared to controls) in the levels of mRNAs encoding pro-tumorigenic factors (*Vegfa*, *S100a8/9*, and *Cd14*) in Sepp1-depeleted tumors (Figure S9d).

Lastly, we evaluated the impact of paquinimod, a S100a9 inhibitor, on tumor growth in sh*Sepp1* and control mice (Figure S9e). Our data showed that paquinimod treatment also significantly attenuated tumor growth in sh*Sepp1* mice compared to controls (Figures S9f-i), These results suggested that targeting neutrophil senescence may be a promising strategy for mitigating Sepp1-depeleted liver cancer.

### Lrp8-mediated uptake of Sepp1 by neutrophils potentiates intracellular hydrogen selenide levels

Prior studies have shown that low-density lipoprotein receptor-related protein 8 (Lrp8), the primary receptor for Sepp1, is displayed on the surfaces of neutrophils^57^. LRP8 has been shown to facilitate the endocytic uptake into cells of LRP8’s ligand, SEPP1, along with associated adaptor proteins^26,28,29^. Building on these insights, we sought to investigate the functional significance of the Sepp1-Lrp8 axis in neutrophil biology, particularly regarding the role of this axis in Sepp1 transport within TINs. To this end, we knockout Lrp8 specifically in neutrophils using *Lrp8*^fl/fl^; *S100a8*^Cre^ (*Lrp8*^ΔNeu^) mice. In line with the findings in tumor -specific Sepp1-depeleted mice, *Lrp8*^ΔNeu^ mice developed significantly increased tumor burden compared to control animals (*S100a8*^Cre^ mice) evidenced by liver-to-body weight ratios and macroscopic tumor counts (Figures 6a-c).

**Figure 6:**
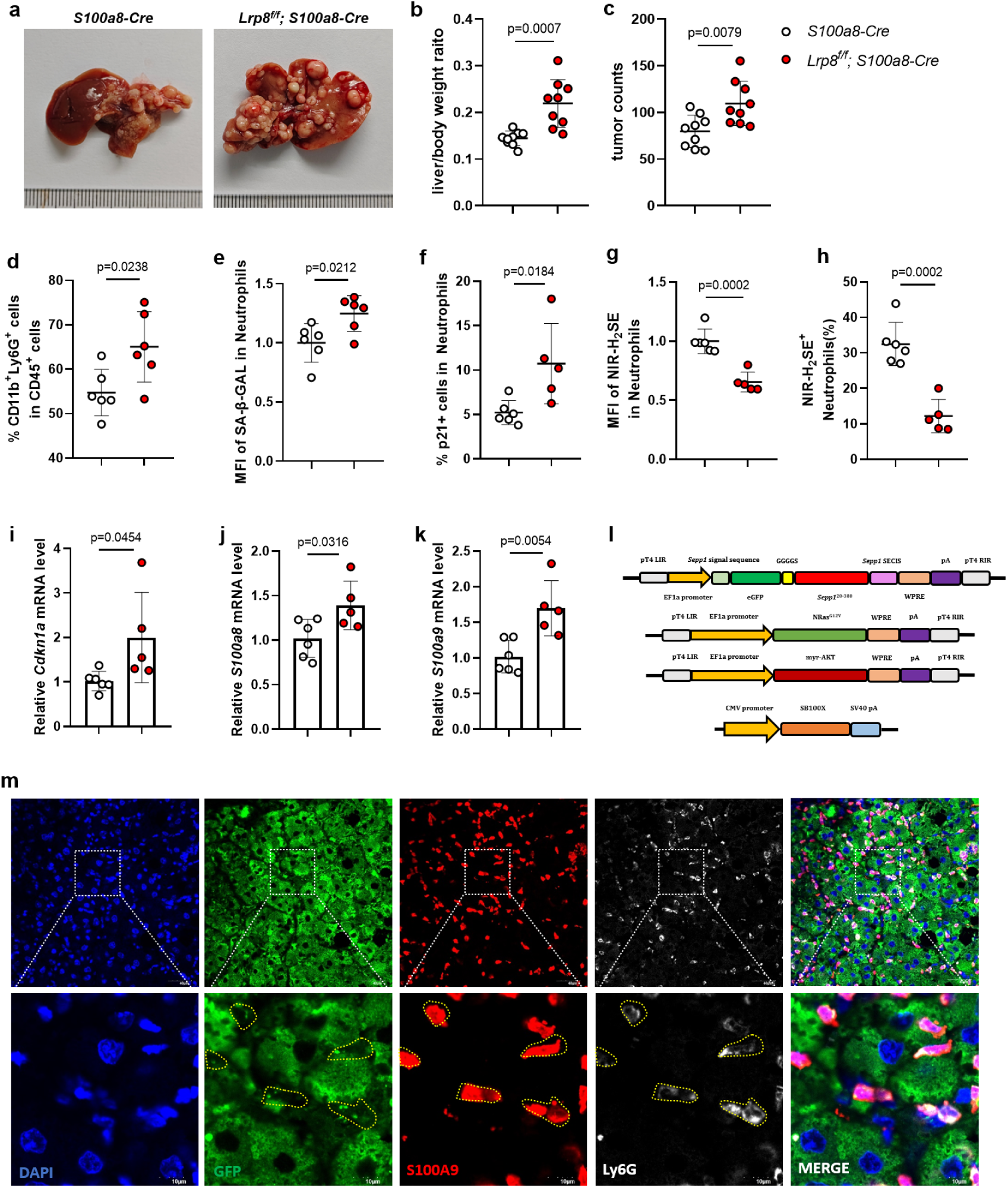
Lrp8-mediated uptake of Sepp1 by neutrophils potentiates intracellular hydrogen selenide levels. (a) Representative images of livers from *S100a8-Cre* and *Lrp8^f/f^; S100a8-Cre* (*Lrp8*^ΔNeu^) mice at endpoint, demonstrating increased tumor burden in *Lrp8*^ΔNeu^ mice. (b, c) Quantification of liver-to-body weight ratio (b) and tumor counts (c) in *Lrp8*^ΔNeu^ versus control groups, showing a significantly higher tumor burden in the *Lrp8*^ΔNeu^ group (n = 9 per group). (d) Flow cytometry analysis showing an increased percentage of neutrophils in the tumor microenvironment of *Lrp8*^ΔNeu^ mice compared to controls (n = 6 per group). (e, f) Flow cytometric analysis of senescence markers in tumor-infiltrating neutrophils from represent mice. Senescence-associated β-galactosidase (SA-β-Gal) positive neutrophils (e) and p21^+^neutrophils (f) were significantly increased in *Lrp8*^ΔNeu^ mice compared to control group (n = 5-6 per group). (g, h) Flow cytometric analysis of neutrophils isolated from *Lrp8*^ΔNeu^ and control tumors. Representative histogram showing NIR-H_2_SE fluorescence, a hydrogen selenide probe, in neutrophils. Quantification of the percentage of NIR-H_2_SE^+^ neutrophils (g) and mean fluorescence intensity (MFI) of NIR-H_2_SE in neutrophils (h), showing significantly reduced hydrogen selenide levels in neutrophils after Lrp8 knockout (n = 5–6 per group). (i) Quantitative RT-PCR analysis of senescence-associated markers *Cdkn1a* in tumor-infiltrating neutrophils (n = 5–6 per group). (j, k) mRNA expression levels of *S100a8* and *S100a9* significantly elevated in tumor-infiltrating neutrophils from *Lrp8*^ΔNeu^ mice (n = 5-6 per group). (l) Schematic representation of hydrodynamic tail vein injection (HDTVi) constructs. Mice were injected with constructs encoding oncogenic NRas^G12V^ and myr-AKT along with a plasmid that encodes a green fluorescent protein (GFP) -Sepp1 fusion protein for tracking Sepp1 localization. (m) Representative immunofluorescence images of liver tissues showing GFP (green) indicating Sepp1 localization, S100A9 (red) indicating neutrophils, lymphocyte antigen 6 family member G (Ly6G; white) as another neutrophil marker, and 4′,6-diamidino-2-phenylindole (DAPI; blue) for nuclei. Images demonstrate the colocalization of GFP with neutrophils, indicating that neutrophils uptake Sepp1. The lower row of images shows magnified views of the boxed regions, highlighting Sepp1 uptake by neutrophils. Data are presented as mean ± SEM; statistical significance was determined using two-tailed unpaired Student’s t-tests.

Flow cytometric analysis identified a significant increase in the percentage of neutrophils in *Lrp8*^ΔNeu^ tumors compared to control tumors (Figure 6d), and a significant upregulation in the proportion of SA-β-Gal positive neutrophils and of p21^+^ neutrophils in the *Lrp8* ^Δ Neu^ tumors compared to control mice (Figures 6e, f). To determine the functional consequences of Sepp1 uptake, we next assessed the functional impact of Sepp1 uptake on intracellular hydrogen selenide levels in neutrophils. Recent advances in selenium biology have shown that intracellular selenium metabolism involves the metabolism and chemical reduction of selenium compounds and selenoproteins to yield hydrogen selenide (HSe^-^/H_2_Se), a pivotal metabolic intermediate^28,29^. We postulated that the absence of Lrp8 or Sepp1 may disrupt selenium homeostasis in neutrophils, impairing the biosynthesis of H_2_Se. Using NIR-H_2_Se, a probe specific for hydrogen selenide detection^58,59^, we conducted flow cytometry on neutrophils isolated from both *Lrp8*^ΔNeu^ and control tumors. Both the proportion of NIR-H_2_Se-positive neutrophils and the MFI were significantly lower in neutrophils from *Lrp8*^ΔNeu^ tumors compared to controls (Figures 6g, h). Moreover, the proportion of NIR-H_2_Se-positive neutrophils and the MFI were also significantly lower in neutrophils from sh*Sepp1* tumors compared to shNC controls, demonstrating that Sepp1 deficiency leads to a decrease in hydrogen selenide levels within neutrophils (Figures S10a-c).

TINs were isolated using MACS, and RT-qPCR analysis revealed significantly increased accumulation, in the *Lrp8^Δ Neu^* group compared to controls, of transcripts encoding senescence markers, including such as *Cdkn1a*, *Cdkn2a* and *Plaur* (Figures 6i, S10d, e). Additionally, we observed significantly elevated levels of transcripts encoding immunosuppressive markers (including *S100a8/9*, *Cd14*, and *Chil3*) and pro-angiogenic markers (*Vegfa*) (Figures 6j, k, S10f-h).

In parallel, *Lrp8* knockout resulted in a significant increase in the percentage of PD-1^+^ CD8^+^ T cells (Figure S10i). Concurrently, we observed decreased expression of GZMB and IFNγ (Figures S10j, k), indicating decreased cytotoxic activity of CD8^+^ T cells in *Lrp8*^ΔNeu^ tumors compared to control tumors. Furthermore, the proportion of NK cells was significantly diminished in the *Lrp8*^ΔNeu^ group compared to the control (Figure S10l), suggesting a broad suppression of cytotoxic immune responses.

To further elucidate the mechanistic link between tumor-derived Sepp1 and neutrophils, we developed a novel experimental system by hydrodynamically injecting mice with constructs encoding oncogenic NRas^G12V^ and myr-AKT, along with a plasmid encoding a Sepp1-enhanced green fluorescent protein (eGFP) fusion protein (permitting precise tracking of Sepp1 localization) (Figure 6l). The ability of neutrophils to uptake Sepp1 was examined using immunofluorescence analysis of tumor tissues. Representative images of the tumors from the injected animals showed that eGFP-labeled Sepp1 (green) colocalized with neutrophil markers S100a9 (red) and lymphocyte antigen 6 family member G (Ly6G; ^29^white); 4′,6-diamidino-2-phenylindole (DAPI; blue) was used to counter-stain the nuclei (Figure 6m). This spatial correlation provided direct visual evidence that TINs actively uptake tumor-derived Sepp1.

These results indicated that Sepp1 is capable of uptake by TINs via Lrp8 and suggests that the protein is essential for maintaining intracellular selenium metabolism.

### Sepp1 regulates neutrophil senescence via H3K4me3

To investigate the factors driving the increased expression of senescence-associated and pro-tumorigenic genes in neutrophils following tumor-specific knockdown of Sepp1, we conducted CUT&RUN sequencing^60,61^ focused on histone modifications and epigenetic changes in TINs. Specifically, this analysis examined the presence on histone H3 protein of trimethylation of lysine 4 (H3K4me3) and acetylation of lysine 27 (H3K27ac), which are classical chromatin marks linked to active gene transcription, and of dimethylation of lysine 9 (H3K9me2), a heterochromatin marker associated with transcriptional repression.

Genome-wide profiling of histone modifications using CUT&RUN sequencing revealed significant enrichment of H3K4me3 marks in chromatin from neutrophils isolated from Sepp1-depeleted tumors compared to those in the chromatin of controls (Figure 7a, left panel). Targeted analysis of H3K4me3 enrichment in 332 senescence-associated genes (identified in senescent-like neutrophil Subcluster 2) demonstrated pronounced hypermethylation at these loci (Figure 7a, right panel). However, the knockdown of Sepp1 in tumors did not significantly alter the levels of H3K27ac or H3K9me2 in the chromatin of neutrophils isolated from Sepp1-depeleted tumors (compared to those in the chromatin of controls) (Figures S11a, b). Gene-specific analysis confirmed substantial increases in H3K4me3 deposition at the promoter regions of key senescence-related genes, including the promoters of *Cdkn1a*, *S100a8*, *Chil3*, and *Vegfa*, in neutrophils from Sepp1-depeleted tumors (Figure 7b). These findings suggested that Sepp1 deficiency induces extensive epigenetic reprogramming, driving neutrophils toward a senescent-like state that promotes tumor progression.

**Figure 7:**
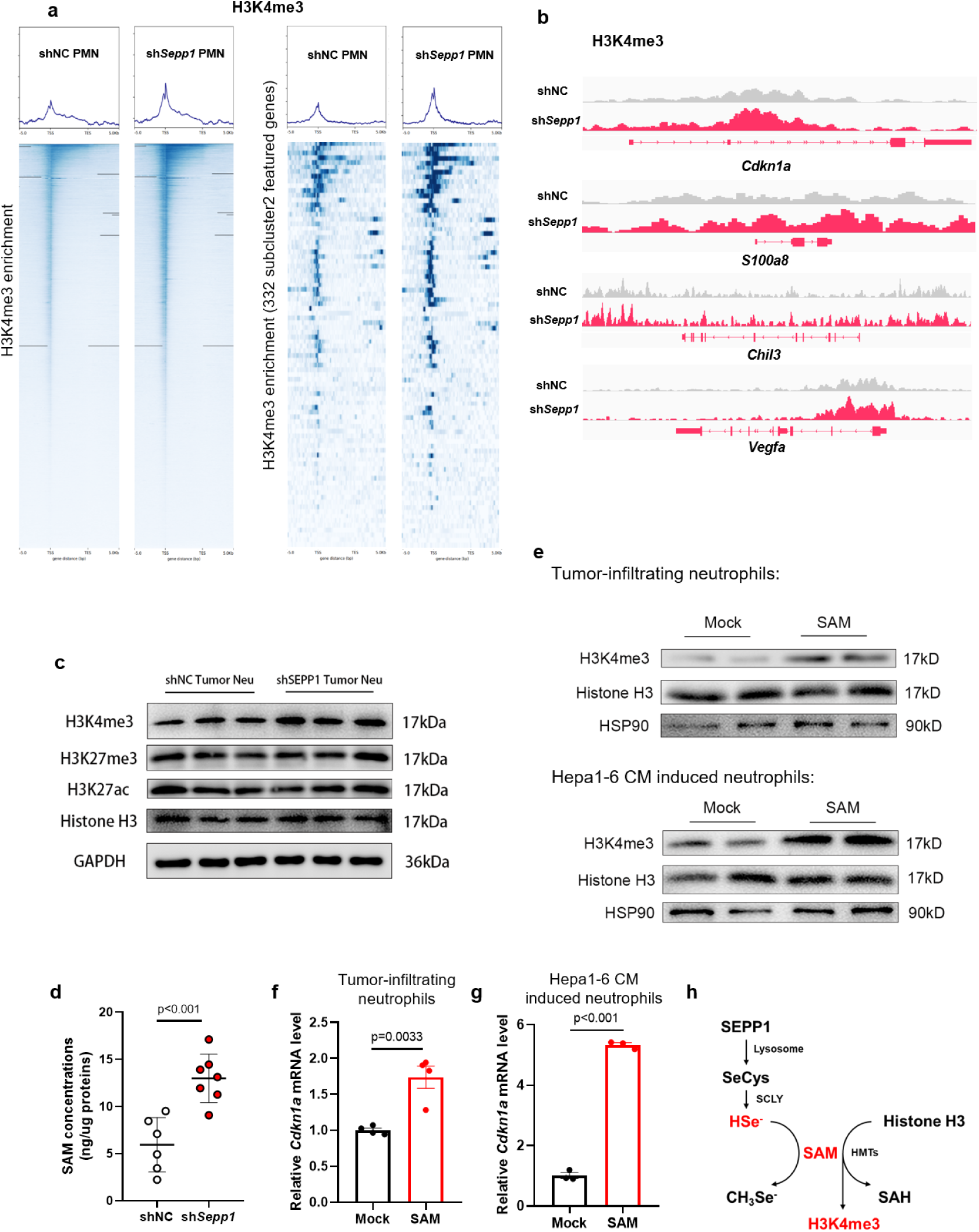
Sepp1 regulates neutrophil H3K4me3 and senescence via hydrogen selenide. (a) Heatmaps and enrichment profiles of trimethylation of lysine 4 on histone 3 (H3K4me3) in neutrophils isolated from shNC and sh*Sepp1* tumors. The left panel shows genome-wide H3K4me3 enrichment profiles in shNC and sh*Sepp1* neutrophils. The right panel highlights H3K4me3 enrichment in the featured genes within the senescent-like subcluster 2, showing increased enrichment in neutrophils from sh*Sepp1* tumors compared to shNC. (b) Genome browser tracks showing H3K4me3 modifications at selected gene loci (*Cdkn1a*, *S100a8*, *Chil3*, *Vegfa*) in neutrophils from shNC and sh*Sepp1* tumors. (c) Western blot analysis of H3K4me3, histone H3, and glyceraldehyde 3-phosphate dehydrogenase (GAPDH; loading control) in tumor-infiltrating neutrophils (TINs; “Neu” in figure) from shNC and sh*Sepp1* tumors. H3K4me3 levels are reduced in sh*Sepp1* neutrophils. Additional histone H3 modifications (acetylation or trimethylation at lysine 27; H3K27ac or H3K27me3, respectively) showed no change. (d) Measurement of S-adenosylmethionine (SAM) concentration in neutrophils, showing increased SAM levels in sh*Sepp1* conditions. (e) Supplementing the culture medium with SAM significantly increased the level of H3K4me3 modification. (f, g) Supplementing the culture medium with SAM significantly increased the expression of *Cdkn1a* in TINs (f) and Hepa1-6-conditioned medium (CM) -induced neutrophils (g). (h) Schematic diagram depicting the proposed mechanism by which Sepp1 regulates H3K4me3 levels via hydrogen selenide (HSe^−^). Data are presented as mean ± SEM; statistical significance was determined using two-tailed unpaired Student’s t-tests.

Western blot analysis confirmed the increase in H3K4me3 levels, but not in H3K27me3 or H3K27ac levels, in neutrophils from Sepp1-depeleted tumors (Figure 7c). In these studies, our data suggested that the absence of Sepp1 deprives neutrophils of selenium substrate, which is required for H_2_Se generation. This deficit would disrupt the H_2_Se/SAM balance. It was well established that SAM is the primary methyl donor for histone methylation; indeed, using ELISA-based quantification, we observed significantly elevated intracellular SAM levels in neutrophils isolated from Sepp1-knockdown tumors compared to control tumors (Figure 7d). To establish the functional consequences of SAM-mediated H3K4me3 modification on neutrophil senescence, we performed in vitro experiments using magnetically sorted TINs. Growth of these cells in culture medium supplemented with SAM significantly enhanced H3K4me3 levels and upregulated *Cdkn1a* expression compared to those grown in cells grown in unsupplemented medium (Figures 7e, f). Other senescence markers, including such as *Cdkn2a*, *Cdkn1b* and *Cdkn2d*, immunosuppressive markers (including *S100a8/9* and *Cd14*) and pro-angiogenic markers (*Vegfa*) are also upregulated (Figure S11c). Similar results were obtained with BM-derived neutrophils stimulated with Hepa1-6 tumor-conditioned medium (TCM) (Figures 7e, g, S11d). To better understand the link between Sepp1 and neutrophil epigenetic regulation, we propose a mechanistic model (Figure 7h) wherein Sepp1 deficiency disrupts selenium metabolism, leading to decreased hydrogen selenide (H₂Se) bioavailability. This metabolic perturbation enhances the conversion of methionine to SAM, resulting in aberrant H3K4me3 accumulation and subsequent transcriptional activation of senescence-associated genes^62^. Our results establish Sepp1 as a critical regulator of neutrophil epigenetics through this protein’s role in maintaining selenium homeostasis and proper SAM metabolism. We believe that this model provides novel insights into the intersection of selenium biology, epigenetic regulation, and tumor immunology.

### SEPP1 expression in human HCC and its correlation with S100A9 levels in clinical HCC samples

Analysis of protein expression levels in the Clinical Proteomic Tumor Analysis Consortium (CPTAC) cohort revealed significantly decreased concentrations of SEPP1 in primary HCC tumors compared to adjacent normal liver tissues (Figure 8a) ^63,64^. Kaplan-Meier survival analysis^65^ showed that patients with higher *SEPP1* mRNA levels exhibited significantly improved overall survival compared to those with lower *SEPP1* expression (Figure 8b). In contrast, elevated transcript levels of *S100A8* and *S100A9* correlated significantly with poorer overall survival in HCC patients (Figures 8c, d). Notably, with the help of the TIMER^66,67^ web tools, we identified a significant inverse correlation between *SEPP1* and *S100A9* mRNA levels (Figure 8e). Next, we analyzed the mRNA levels of *SEPP1*, *S100A8*, and *S100A9* in non-tumor (NT) and tumor (T) tissues from a cohort of patients seen at the Changhai Hospital. The results obtained from the analysis of this cohort of patients were consistent with those obtained with the database (Figure 8f, g, h). Specifically, *SEPP1* mRNA expression was significantly decreased in tumor tissues from Changhai Hospital patients (compared to matched non-tumor liver tissues). Conversely, *S100A8* and *S100A9* expression was significantly higher in HCC tissues compared to matched non-tumor liver samples (Figures 8g, h).

**Figure 8:**
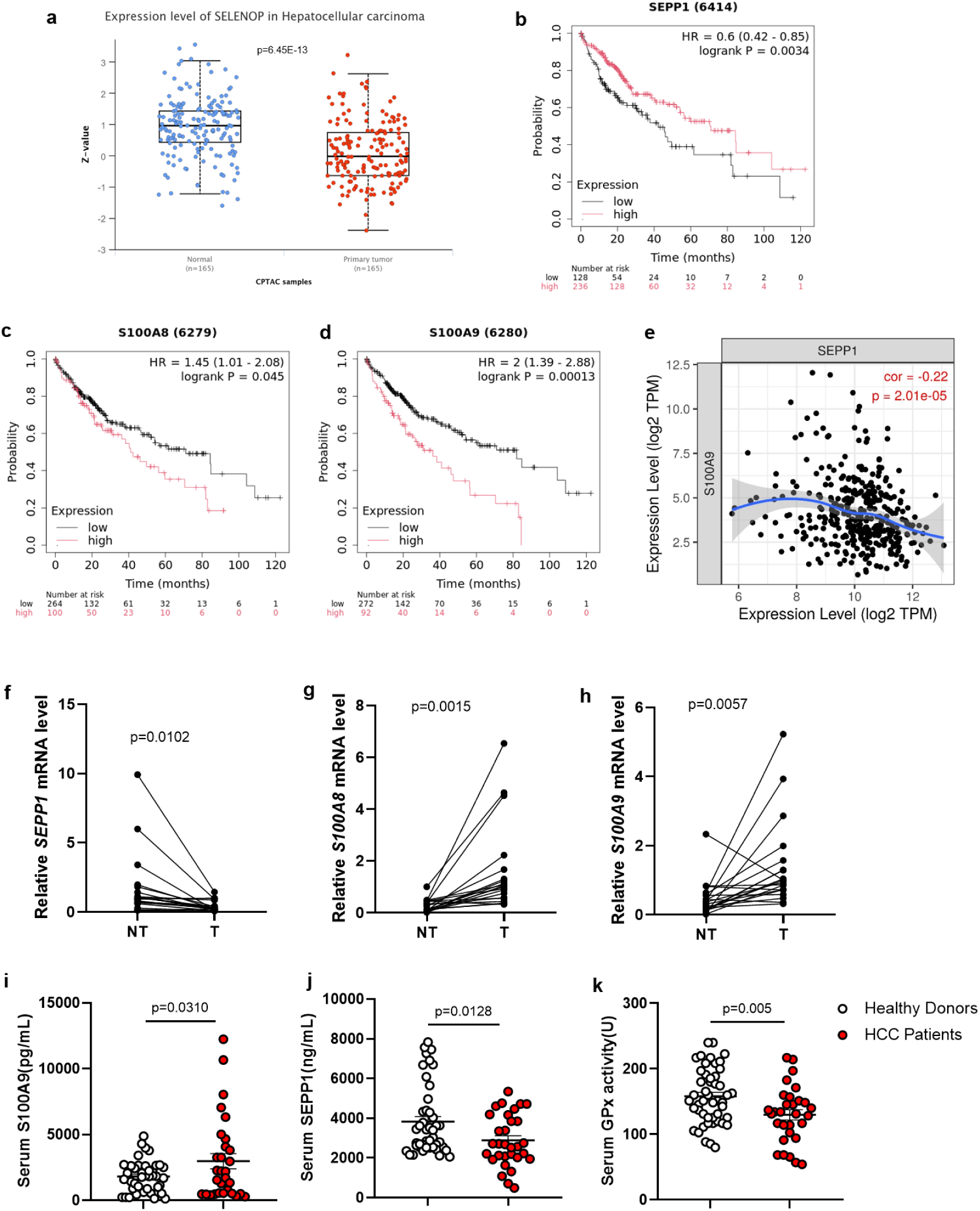
SEPP1 expression in human HCC and correlation of SEPP1 levels with those of S100a9. (a) Box plot of SEPP1 (SELENOP) protein levels in hepatocellular carcinoma (HCC) tissues compared to normal liver tissues, based on Clinical Proteomic Tumor Analysis Consortium (CPTAC) samples (n = 165 for HCC, n = 168 for normal). (b) Kaplan-Meier survival analysis of HCC patients stratified by SEPP1 levels (HR = 0.6, p = 0.0034). (c, d) Kaplan-Meier survival analyses of HCC patients stratified by S100A8 (c) and S100A9 (d) levels. Higher levels of S100A8 and S100A9 were associated with poorer survival outcomes (HR = 1.45, p = 0.045 for S100A8; HR = 2.0, p = 0.00013 for S100A9). (e) Scatter plot showing the negative correlation between SEPP1 and S100A9 levels in HCC tissues (correlation coefficient = -0.22, p = 2.01E-05), suggesting an inverse relationship between SEPP1 and S100A9 levels in HCC. (f–h) Paired analysis of mRNA expression in HCC tissues (T) versus non-tumor adjacent tissues (NT) from HCC patients. (f) *SEPP1* mRNA levels were significantly decreased in tumor tissues compared to non-tumor tissues. (g, h) *S100A8* (g) and *S100A9* (h) mRNA levels are significantly elevated in tumor tissues compared to non-tumor tissues (n=20). (i–k) Serum analysis comparing healthy donors and HCC patients. (i) Serum S100A9 protein levels were significantly higher in HCC patients (n=30) compared to healthy donors (n=42). (j) Serum SEPP1protein levels were significantly lower in HCC patients (n=31) compared to healthy donors (n=44). (k) Serum glutathione peroxidase (GPx) activity is decreased in HCC patients (n=30) compared to healthy donors (n=48). Data are presented as mean ± SEM; statistical significance was determined using two-tailed unpaired Student’s t-test.

To establish the translational relevance of these findings, we measured circulating biomarkers in serum samples from HCC patients and healthy controls seen at Changhai Hospital. HCC patients exhibited significantly elevated serum S100A9 levels (Figure 8i) along with decreased serum SEPP1 concentrations (Figure 8j) and decreased serum glutathione peroxidase (GPx) activity compared to healthy donors (Figure 8k).

These clinical findings collectively demonstrated that a decrease in SEPP1 levels is a hallmark of HCC progression, and constitutes a parameter that is associated with poor prognosis and inversely correlated with S100A9 expression. Serum biomarker analysis further confirmed decreased SEPP1 and GPx activity and elevated S100A9 levels in HCC patients compared to healthy controls, highlighting the potential utility of these biomarkers as diagnostic and prognostic indicators in HCC. Our study elucidated a novel mechanism by which selenoprotein P governs the immunosuppressive TME in HCC through regulation of neutrophil biology. We demonstrated that SEPP1 deficiency drives HCC progression by promoting the expansion of a senescent-like neutrophil population characterized by immunosuppressive properties and pro-tumorigenic functions. These findings establish SEPP1 as a critical regulator of the hepatic TME and provide new insights into the role of selenium metabolism in innate immunity and cancer progression.

## Discussion

Our study unveils a previously unrecognized immunometabolism axis centered on selenium homeostasis, an axis that orchestrates neutrophil senescence and shapes the selenium deficiency-associated immunosuppressive TME in HCC. Through the analysis of single-cell RNA sequencing data, we showed that this transformational subtype of neutrophil is characterized by the accumulation of transcripts encoding canonical senescence markers (e.g., Cdkn1a), immunosuppressive effectors (S100a8/9, Chil3, and Cd14), and-angiogenic mediators (e.g., Vegfa). Furthermore, we demonstrated that Sepp1 deficiency blocks selenium metabolism by the Sepp1/hydrogen selenide/SAM pathway in these TINs, and showed that the increase in SAM level leads to a profound epigenetic rewiring that drives neutrophils into a distinct senescence-like state. Importantly, we showed that S100a8/9 serves as a causative molecule with immunosuppressive properties, operating downstream of this selenium deficiency-mediated senescence program, creating a feed-forward loop that potentiates the immunosuppressive microenvironment and worsened prognosis. We believe that these findings not only redefine our understanding of myeloid cell biology in the TME but also reveal a targetable selenium metabolic vulnerability; focusing on those neutrophils may be a promising therapeutic strategy in HCC.

Multiple distinct developmental states of neutrophils co-exist within the TME^68–71^. Accumulating clinical evidence has consistently demonstrated that elevated infiltration by TANs correlates significantly with enhanced tumor progression, increased metastasis and unfavorable clinical outcomes across various solid malignancies^54,55^. Particularly in HCC, TANs have emerged as a critical prognostic indicator, with recent studies validating their utility as a robust biomarker for disease progression^72^. Across diverse murine models of HCC, neutrophil infiltration is consistently observed, yet the relative abundance of neutrophils varies markedly. In our HCC mouse models, neutrophils constitute the predominant immune cell population within TMEs. In contrast, we found that classical carcinogen-induced models such as DEN-driven HCC demonstrate comparatively limited neutrophil recruitment. This disparity is supported by single-cell RNA sequencing studies of DEN-induced HCC^73,74^, which have corroborated the relatively low neutrophil prevalence in these systems^75^. The inherent technical challenges associated with single-cell RNA sequencing, particularly in capturing short-lived neutrophils, have recently been overcome, enabling comprehensive characterization of TAN heterogeneity. Pioneering work from the Zhang laboratory, employing large-scale single-cell transcriptomic profiling of liver cancer specimens, has identified previously unrecognized pro-tumorigenic neutrophil subsets, thereby suggesting that neutrophils might serve as therapeutic targets in cancer immunotherapy^43,50,71^.

Immunosuppressive neutrophils have been shown to contribute to tumorigenesis across various cancer types through diverse mechanisms^68,76–78^. In recent years, immunosuppressive TINs have been identified as key players in promoting therapy resistance in HCC^50^. These findings have driven efforts to develop novel therapeutic strategies aimed at blocking the recruitment of these cells to the TME. For example, several drugs targeting immunosuppressive neutrophils have demonstrated enhanced antitumor efficacy when combined with immune checkpoint inhibitors such as anti-PD-1 antibodies. By blocking the recruitment of polymorphonuclear myeloid-derived suppressor cells (PMN-MDSCs), C-X-C chemokine receptor 2 (CXCR2) inhibitors impede tumor growth and exhibit synergistic effects with anti-PD-1 antibodies^79,80^. Fatty acid transport protein 2 (FATP2) inhibitors, which regulate the immunosuppressive functions of immunosuppressive neutrophils, also attenuate tumor growth and potentiate the effects of anti-cytotoxic T-lymphocyte associated protein 4 (anti-CTLA4) antibodies^81^. Ferroptosis inhibitors like liproxstatin-1 diminish the immunosuppressive capacity of neutrophils by inhibiting neutrophil ferroptosis, thereby enhancing the efficacy of anti-PD-1 therapy^82^. Additionally, inhibiting CD300ld on the surface of neutrophils has been shown to improve the antitumor efficacy of anti-PD-1 treatment^3^. Indeed, it previously has been reported that immunosuppressive neutrophils express markers of senescence in the prostate TME, and that the histone deacetylase (HDAC) inhibitor romidepsin improves the efficacy of cancer therapy efficacy, presumably by eliminating senescent neutrophils^11^. We therefore wondered whether, in HCC, the senescence characteristics of neutrophils may be a central regulatory factor underlying the immunosuppressive and tumor-promoting functions of this class of cells.

Selenium, an essential trace element, plays a pivotal role in maintaining cellular homeostasis and overall health. However, given the complexity of the intracellular metabolism of selenium compounds, this metabolic pathway remains poorly understood^83–85^. In the present study, we provide compelling evidence that TINs can uptake selenoprotein P, as demonstrated by the colocalization of eGFP-labeled Sepp1 with neutrophil-specific markers. Our findings identify Sepp1 as a critical extrinsic selenium reservoir for neutrophils. Previous studies have indicated that selenium compounds or selenoproteins are metabolized to hydrogen selenide (HSe^-^/H_2_Se), a key intermediate in selenium metabolism. We hypothesize that the absence of Sepp1 deprives neutrophils of the selenium substrate necessary for H_2_Se generation, thereby disrupting the H_2_Se/SAM balance. H_2_Se, by functioning as a methyltransferase inhibitor, normally modulates intracellular SAM levels^62,86,87^. SAM, a universal methyl donor, is essential for the methylation of diverse biomolecules, including DNA, RNA, proteins, and neurotransmitters^88,89^. In the present study, we demonstrated that the deficiency of H_2_Se leads to uncontrolled SAM accumulation, resulting in hypermethylation of histones, as evidenced by elevated H3K4me3 modifications, and a marked accumulation of the *Cdkn1a* transcript in neutrophils. This epigenetic reprogramming drives neutrophils into a senescence-associated immunosuppressive state, mediated by the Sepp1/H_2_Se/SAM axis. The central role of Sepp1 metabolism in regulating neutrophil senescence underscores this protein’s potential as a therapeutic target in tumor immunology, warranting further investigation into its broader implications in cancer progression and immune evasion^90,91^.

Cellular senescence, a state of stable exit from the cell cycle, often has been compared to apoptosis as an intrinsic mechanism of tumor suppression^92^. Senescence has been assumed to be functionally similar to apoptotic cell death, in terms of its effects on tumor suppression^93,94^. However, this paradigm has been increasingly challenged by recent studies revealing the dual roles of senescence in tumor biology. Various triggers induce either senescence or apoptosis, depending on the cell-type tissue environment and the nature of the inducing stimulus. As senescent cells are permanently arrested, the aging process of immune cells profoundly compromises immune surveillance and contributes to tumor progression^95,96^. In the present study, we demonstrated that Sepp1 deficiency induces a profound metabolic shift, characterized by elevated SAM levels, which drives H3K4me3-mediated epigenetic reprogramming. This reprogramming upregulates the expression of *Cdkn1a*, a key senescence marker, and promotes the SASP in neutrophils. The SASP also has been reported to include proteins that can help senescent cells evade immune recognition and clearance. We further identified Sepp1 as a novel regulator of neutrophil senescence and showed that Sepp1-depeleted senescent neutrophils exacerbate T cell exhaustion while impairing (both in vitro and in vivo) the function of cytotoxic CD8^+^ T cells. These findings highlight a previously unrecognized role for senescent neutrophils in fostering an immunosuppressive TME, thereby potentiating tumor progression.

Our findings revealed that TINs uptake Sepp1 via Lrp8 and transport the protein into the cell. Previous studies have demonstrated that Lrp8, the primary receptor for Sepp1, is expressed on the surface of neutrophils^57^. Lrp8 has been shown to mediate the endocytic uptake of Sepp1 into cells, along with its associated adaptor proteins^26,28,29^. Notably, Leiter et al. demonstrated that selenium supplementation reverses cognitive deficits induced by hippocampal injury and aging, and further showed that Sepp1 and its receptor Lrp8 are indispensable for exercise-induced adult neurogenesis^97^. Building on these insights, we genetically ablated Lrp8 specifically in neutrophils and observed a marked alteration in their functional phenotype. Notably, Lrp8 knockout significantly attenuated neutrophil senescence within the tumor microenvironment, accompanied by a substantial decrease in intracellular hydrogen selenide (H₂Se) levels. These results indicate that Lrp8-mediated Sepp1 uptake is a critical determinant of selenium metabolism in neutrophils and directly influences their function in the tumor context. These investigations are expected to yield critical insights into the interplay between metabolic regulation and immune modulation in cancer progression.

From a clinical perspective, our findings significantly advance the understanding of the TME by elucidating the pivotal role of selenium status and SEPP1 function in modulating antitumor immunity. Notably, in the present work, selenium supplementation restored SEPP1 expression and attenuated tumor progression in our experimental models; this supplementation also demonstrated a synergistic effect with immune checkpoint blockade, markedly enhancing the efficacy of anti-PD-1 therapy. This compelling synergy underscores the potential utility of correcting selenium imbalances as a low-toxicity, clinically feasible strategy to augment the response to immunotherapy, which remains suboptimal and highly variable across patient populations.

Our study also highlights critical avenues for future research. The molecular mechanisms underlying the interplay between selenium deficiency and epigenetic reprogramming of immune cells warrant further investigation, particularly in the context of how other micronutrients and selenoproteins may contribute to immune cell modulation. Moreover, while our preclinical findings are robust, translational clinical studies are imperative to define optimal dosing regimens, therapeutic timing, and patient stratification strategies for the use of selenium supplementation as an adjuvant to immunotherapy. Identifying predictive biomarkers—both tumor-intrinsic and host-derived—will be crucial to delineate which patients and tumor types are most likely to derive therapeutic benefit from this intervention. We propose that such precision medicine approaches may pave the way for integrating selenium-based strategies into the evolving landscape of combination cancer therapies.

In sum, our study elucidates the pivotal roles of selenium and SEPP1 as master regulators of the immune TME in HCC. We employed single-cell RNA sequencing in Sepp1-depeleted murine HCC models to identify a novel senescence-associated neutrophil subpopulation co-expressing *Cdkn1a*, *S100a8/9*, and *Vegfa*. This subset drives immunosuppression and tumor progression. Mechanistically, the knockdown of tumor *Sepp1* disrupts the transport of the corresponding protein as well as metabolic pathways in TINs, leading to H_2_Se depletion, SAM accumulation, and enhanced H3K4me3 histone modifications. These changes collectively foster a pro-senescence chromatin environment in TINs. Additionally, we showed that selenium supplementation not only restores Sepp1 expression but also plays a crucial role in reshaping anti-tumor immunity by reversing the senescence-like phenotype and synergizing with anti-PD-1 therapy. Our findings highlight SEPP1 as a critical regulator countering neutrophil senescence and its immunosuppressive effects in HCC. We also explored the therapeutic potential of targeting senescent-like neutrophils and the role of selenium supplementation as an adjuvant to enhance immunotherapy efficacy in liver cancer. This paradigm, once clinically validated, may provide an important step forward in personalized cancer treatment, broadening the arsenal of tools available to clinicians seeking to enhance antitumor immunity.

## Methods

### Mice

All animals were maintained and used in accordance with the guidelines of, and under approval by, the Institutional Animal Care and Use Committee of the Shanghai Institute for Nutrition and Health (Ethical Committee Approval No. SINH-2024-ZLX-1). Only healthy male C57BL/6J mice were used for the experiments described here. Animals were purchased from the Shanghai Laboratory Animals Center (SLAC) of the Chinese Academy of Sciences. Lrp8^flox/flox^ mice and S100a8^Cre^ mice were purchased from Cyagen (Guangzhou, China). OT-I TCR-transgenic mice were provided by Dr. Ju Qiu (Professor, Laboratory of Inflammation and Immune Regulation, Shanghai Institute of Nutrition and Health). *Cas9* knock-in mice (*Rosa26^Cas9+/+^*) were provided by Qiurong Ding (Professor, Laboratory of Stem Cell and Liver Disease, Shanghai Institute of Nutrition and Health. Mice were maintained in a temperature-controlled room with 12-h/12-h light/dark diurnal cycle. Throughout the study, animals were provided with *ad libitum* access to food and water.

### Plasmids

Using the pT4 backbone (Addgene, #108352), a series of recombinant plasmids were constructed in which specific oncogenes are driven by the *EF1a* promoter. For this purpose, the target gene fragments were amplified and subsequently inserted downstream of the *EF1a* promoter using Gibson assembly^98^. The target oncogenes inserted into the recombinant plasmids included loci encoding NRas^G12V^, myr-AKT, c-Myc^T58A^, Yap^S127A^, and ΔN90β-catenin.

To construct the pT4-shRNA plasmids, shRNA targeting *Sepp1*(sh*Sepp1*-1: GCTCCTGTGTAAGTTGTCTAA, sh*Sepp1*-2: GCTGAGATAATCAGACCAGGA) or control (shNC-1: CCTAAGGTTAAGTCGCCCTCG, shNC-2: GTTCAGATGTGCGGCGAGT) sequences were initially cloned (using PCR amplification) into the pLKO.1 vector (Addgene, #8453) to generate the U6-shRNA expression cassette. The pT4-NRas^G12V^ plasmid was linearized by HindIII single digestion and dephosphorylated. Subsequently, two distinct U6-shRNA expression cassettes were ligated into the linearized pT4-NRas^G12V^ plasmid using Gibson assembly.

For the construction of the expression plasmid encoding the eGFP-Sepp1 fusion protein, the full-length coding sequence (CDS) and 3′-untranslated region (UTR) sequences of *Sepp1* were amplified from mouse liver cDNA and cloned into the pT4 vector. The resulting pT4-Sepp1 vector then was linearized by PCR, and the *eGFP* CDS was inserted (using Gibson assembly) immediately downstream of (and in-frame with) the DNA sequences encoding the Sepp1 signal peptide, yielding the pT4-eGFP-Sepp1 plasmid.

### Liver tumor mouse model by hydrodynamic tail vein injection (HDTVi)

For all animal studies, laboratory personnel were not blinded to the identities of the test article or dosing groups.

DNA for HDTVi was prepared using Endofree Maxi Plasmid Kit (TIANGEN, DP117). On the day of dosing, 30 μg of the appropriate pT4 vectors and 3 ug of the SB100X transposase-encoding plasmid (Addgene, #34879) were diluted in sterile filtered Ringer’s solution to a total volume of 10% of the body weight of the 8-week-old animal; the resulting dosing solution then was injected into a mouse lateral tail vein as a bolus injection lasting less than 7 s.

For the selenium supplementation experiment, sodium selenite or selenomethionine (Sigma-Aldrich, S3132) was added at 10 ppm to the drinking water of mice starting 14 days after HDTVi. The mice were allowed free access to the supplemented water.

For selenium and immune checkpoint inhibitor treatment, mice were randomized and assigned to control or treatment groups at 14 days after HDTVi. The groups received twice-weekly intraperitoneal (IP) injections of (respectively) isotype antibody (2A3, Bioxcell) or anti-PD-1 antibody (RMP1-14, Bioxcell) at 200 μg/mouse. Where applicable, selenomethionine was administered five times per week via IP injection at 2 mg/kg body weight; control animals received the equivalent volume (0.1 mL/mouse) of PBS, the vehicle used to formulate selenomethionine.

For neutrophil depletion, mice were randomized and assigned to control or treatment groups at 14 days after HDTVi. The groups received IP injection of (respectively) isotype antibody (2A3, Bioxcell) or anti-Ly6G antibody (1A8, Bioxcell) at 100 μg/mouse on every other day (Q2D).

For senolytic treatment, dasatinib hydrochloride (D; MedChem Express, HY-10181A) and quercetin (Q; MedChem Express, HY-18085) were dissolved in dimethyl sulfoxide (DMSO) to respective concentrations of 15 and 30 mg/mL. This DMSO stock then was diluted in an aqueous vehicle to generate a dosing solution containing dasatinib and quercetin at final concentrations of

1.25 and 12.5 mg/mL (respectively) in a mixture consisting of 40% PEG300, 5% Tween-80, and 45% saline. As above, mice were randomized and assigned to treatment or control groups at 14 days after HDTVi. The treatment group then was administered twice weekly by IP injection with the D&Q combination at 5 and 50 mg/kg (respectively); the control groups was administered (at the same frequency and by the same route) with the PEG300/Tween 80/saline vehicle (without compounds).

For paquinimod treatment, this reagent (Selleck, S9963) was dissolved in DMSO, then diluted into the PEG300/Tween 80/saline vehicle (as above) to a concentration of 1.25 mg/mL. As above, mice were randomized and assigned to treatment or control groups at 14 days after HDTVi. The treatment group then was administered once daily (QD) by IP injection with 5 mg/kg paquinimod; the control groups was administered (at the same frequency and by the same route) with the PEG300/Tween 80/saline vehicle (without compound).

### Liver tumor mouse model with AAV injections

Liver-specific *Trp53* and *Pten* knockout was achieved using *Cas9* knock-in (*Rosa26^Cas9+/+^*) mice. Adeno-associated virus serotype 8 (AAV8) encoding iCre recombinase and carrying sgRNAs targeting *Tp53* (GCCTCGAGCTCCCTCTGAGCC) and *Pten* (GAGATCGTTAGCAGAAACAAA) were administered by intravenous injection (IV; into a lateral tail vein) at a dose of 2 × 10¹¹ viral genomes (vg) per mouse in volume of 250 µL phosphate-buffered saline (PBS).

For liver-specific *Sepp1* knockout, the constructs consisted of an AAV8-iCre vector carrying sgRNA sequences targeting *Sepp1* (sg*Sepp1*-1 (GCTTGTTACAAAGCCCCGGAG) and sg*Sepp1*-2 (GCAGAAGACACAAGTATCAGC)) or control (sgNC-1 (GACGGAGGCTAAGCGTCGCAA) and sgNC-2 (GACGTGTAAGGCGAACGCCTT)). These AAV8 constructs were injected as above.

### Cell lines

A liver-specific *Trp53* knockout was achieved using *Cas9* knock-in (*Rosa26^Cas9+/+^*) mice. An AAV8 vector encoding iCre recombinase and carrying a sgRNA targeting *Tp53* (sequence: GCCTCGAGCTCCCTCTGAGCC) was injected IV (as above) into 6- to 8-week-old *Cas9* knock-in mice. At 14 days after IV injection, primary hepatocytes were isolated from the resulting *Trp53*-knockout mice (as well as from control wild-type (WT) animals) using a two-step collagenase perfusion method. Briefly, mice were anesthetized with 2,2,2-tribromoethanol, and the liver was perfused (via the portal vein, using a peristaltic pump) with 25 mL perfusion buffer (1× PBS, 0.9 g/L glucose, 0.35 g/L NaHCO_3_, 190 mg/L ethylene-bis(oxyethylenenitrilo)tetraacetic acid (EGTA), pH 7.4) for 3 minutes, followed by 50 mL digestion buffer (1× PBS, 0.2 mg/mL collagenase I (Worthington, LS004196), 15 mM N-2-hydroxyethylpiperazine-N-2-ethane sulfonic acid (HEPES), 0.35 g/L NaHCO_3_, 0.5 mM CaCl_2_, pH 7.4) for 5 minutes. The liver tissue then was excised and filtered through a 70-µm cell strainer. Cell suspensions were centrifuged at 4 ℃, 50*g* for 2 minutes to remove debris. Dead cells were removed by centrifugation in 50% Percoll solution at 4 ℃, 50x *g* for 10 minutes, and the resulting viable hepatocytes were resuspended for further use.

WT and *Trp53*-knockout primary hepatocytes were seeded in 6-cm cell culture dishes and transfected with the combination of representative pT4 oncogene plasmids and the pCMV-SB100X plasmid (5 and 1 µg/well, respectively) using jetOPTIMUS® DNA Transfection Reagent (Polyplus, 101000006) according to the manufacturer’s protocol. Non-transfected hepatocytes, which are non-proliferative, subsequently die, leaving only transfected cells to proliferate and survive for subsequent passage.

The AML12, H22, and Hepa1-6 cell lines were procured from the American Type Culture Collection (ATCC; Manassas, Virginia).

All cell lines, with the exception of AML12 and H22, were cultured in Dulbecco’s Modified Eagle’s Medium (DMEM) supplemented with 10% fetal bovine serum (FBS) and 1% of a 100x penicillin/streptomycin stock. AML12 was cultured in DMEM/F-12 supplemented with 10% FBS, 10 µg/mL insulin, 5.5 µg/mL transferrin, and 1× penicillin/streptomycin. H22 was cultured in RPMI-1640 medium supplemented with 10% FBS and 1× penicillin/streptomycin. All cultured cells were grown at 37 ℃ in a 5% CO_2_ environment

### Flow cytometry

Tumor-bearing mice were euthanized, and liver tumor tissues were isolated, washed with ice-cold PBS to remove blood, and digested using the Tumor Dissociation Kit (Miltenyi Biotec, 130-096-730) according to the manufacturer’s instructions. The tissues were incubated at 37 °C for 30 minutes on a shaker, and the resulting digested mixtures cells were filtered through a 40-µm nylon cell strainer.

For surface staining, cells were incubated with anti-mouse CD16/CD32 monoclonal antibody (eBioscience, 93, 1:200) to block Fc receptor binding. Viability was assessed using the LIVE/DEAD™ Fixable Dead Cell Stain Kit (Thermo Fisher Scientific, L10119) according to the manufacturer’s instructions. Following staining for viability, cells were washed with ice-cold PBS and resuspended in staining buffer. followed by staining with antibodies against CD45 (30-F11, 1:200), CD11b (M1/70, 1:200), Ly6G (1A8, 1:200), CD3 (145-2C11, 1:200), CD4 (GK1.5, 1:200), CD8 (53-6.7, 1:200), NK1.1(PK136,1:200) and CD279(J43, 1:200).

For intracellular staining of cytokines (IFNγ, GZMB, and tumor necrosis factor alpha (TNFα)), single-cell suspensions were prepared in RPMI-1640 medium containing 10% FBS, then adjusted to densities of 1×10^6^ cells/mL. Cells were stimulated by incubation (at 37 °C in 5% CO₂ for 4 hours) in the presence of 50 ng/mL phorbol 12-myristate 13-acetate (PMA) and 1 µg/mL ionomycin; 5 µg/mL brefeldin A also was included to block cytokine secretion. Stimulated cells were washed with PBS, followed by cell viability assessment, Fc receptor blocking, and cell surface staining. Intracellular staining was performed using the BD Cytofix/Cytoperm™ Fixation/Permeabilization Kit (BD, 554714) according to the manufacturer’ s protocol. Intracellular staining employed antibodies against IFNγ (XMG1.2, 1:200) and GZMB (16G6, 1:200).

For intracellular staining of p21 and S100a9, cells again were subjected to Fc receptor block, cell viability assessment, and cell surface staining, in this case followed by intracellular staining using the eBioscience™ Foxp3 / Transcription Factor Staining Buffer Set (eBioscience, 00-5523-00) according to the manufacturer’s protocol. Intracellular staining employed antibodies against p21 (Abcam, ab188224, 1:200) and S100A8/9 (Abcam, ab288715, 1:200) with overnight incubation at 4 °C, followed by staining with secondary antibody goat anti-rabbit IgG (H+L) conjugated to AlexaFluor 568 (Thermo Fisher Scientific, A-11011, 1:200). The samples were analyzed on a Cytoflex LX. Data analysis was conducted using Flow Jo (v10.8.1) software.

### Isolation of neutrophils

For BM neutrophil isolation, single-cell suspensions were prepared from BM. Neutrophils then were isolated by using the EasySep Mouse Neutrophil Enrichment Kit (Stem Cell, 19762) according to the manufacturer’s instructions.

For isolation of TINs, single-cell suspensions of tumor tissues were incubated at 4 °C for 15 min with a rat anti-mouse CD16/CD32 monoclonal antibody (1:100). Biotinylated anti-mouse Ly6G antibody (1:100, BioLegend, 1A8) then was added to each tube, and the mixture was incubated at 4 °C for another 30 min. Finally, MojoSort™ Streptavidin Nanobeads (BioLegend, Cat. No. 480016) were added to each tube, and the mixture was incubated, with gentle mixing, at 4 °C for a further 20 minutes. Labeled cells were isolated using a magnetic separation rack, washed twice with PBS, and resuspended for downstream analysis.

### In vitro culture of neutrophils

Spent medium was collected from cultures of representative cell lines and centrifuged at 1500x *g* at 4 °C for 10 minutes to pellet dead cells and debris. The resulting supernatant then was passed through a 0.45-µm filter to remove any remaining cells and particulates, and the resulting filtrate was aliquoted and stored at −80 °C pending use as TCM. For those experiments, neutrophils (5×10⁵ per well) were cultured in normal medium or TCM at 37 °C in a 5% CO₂ environment. After 24 hours, the cultured neutrophils were used for downstream analysis.

For SAM treatment, MACS was used to isolate liver TINs from animals injected with NRas^G12V^+ myr-AKT-encoding constructs. The isolated cells then were cultured for 24 hours in RPMI-1640 medium supplemented with 50 μM SAM (MedChem Express, HY-B0617A). For SAM treatment of BM neutrophils, the cells were cultured for 24 hours in Hepa1-6 TCM supplemented with 50 μM SAM.

### T cell-suppression assay

TINs were plated in U-bottom 96-well plates in RPMI-1640 medium supplemented with 10% FBS and 0.1 ng/mL OVA_254–267_ peptide (sequence SIINFEKL; InvivoGen, vac-sin) and co-cultured at various ratios with CFSE (Invitrogen, C34554) -labeled splenocytes (2 × 10^5^ OT-1 splenocytes/well) derived from OT-1 mice. After 48 hours of culture, cells were stained for cell viability assessment and also subjected to cell surface staining for CD3 and CD8. Proliferation of CD8⁺ T cells was assessed by flow cytometry. Additionally, the supernatant from the co-culture was collected and stored at −80 °C pending subsequent ELISA analysis.

### Measurement of SA-β-Gal activity by flow cytometry

SA-β-Gal activity was measured as described previously^51^. Briefly, cells were pre-treated by incubation at 37 °C for 1 h in fresh cell culture medium supplemented with 100 nM bafilomycin A1 (MCE. Cat. HY-100558), thereby inducing lysosomal alkalinization. Thereafter, the fluorogenic substrate Xite™ Red β-D-galactopyranoside (AAT Bioquest, Cat. 14035) was added to the cell culture medium to a final concentration of 10 μM. After incubation at 37 °C for 1 h, the cells were used to perform cell viability assessment and cell surface staining. The resulting samples were subjected to flow cytometric analysis on a Cytoflex LX using the phycoerythrin (PE) channel to detect SA-β-Gal.

### Measurement of hydrogen selenide levels by flow cytometry

Tumor tissues were enzymatically digested into single-cell suspensions. The cells then were incubated with 10 μM NIR-H_2_SE probe at 37 °C for 1 hour. After incubation, the cells were used to perform cell viability assessment and cell surface staining for CD45, CD11b, and Ly6G. The resulting samples were subjected to flow cytometric analysis on a Cytoflex LX.

### RNA isolation and RT-qPCR

Total RNA was extracted from tissues or cells using the TRIzol reagent (Thermo Fisher Scientific, Cat. 15596018) according to the manufacturer’s instructions. Reverse transcription of isolated RNA was performed using the Prime Script reverse transcription kit (Takara, Cat. RR047A). Quantitative real-time PCR was carried out using Hieff® qPCR SYBR Green Master Mix (Low Rox Plus) (Yeasen Biotechnology, Cat. 11202ES03) on a Quant Studio 6 machine. Transcript levels were normalized to those of house-keeping transcripts encoding actin or glyceraldehyde 3-phosphate dehydrogenase (GAPDH) in the respective sample. All primer pairs were designed for the same cycling conditions: 5 min at 95 °C for initial denaturation, 40 cycles of 10 s at 95 °C for denaturation, and 30 s at 60 °C for annealing and extension.

### ELISA

Mice were anesthetized with 2,2,2-tribromoethanol and subjected to terminal cardiocentesis. The resulting whole blood (no anticoagulant) was allowed to clot on ice and then centrifuged at 3000x *g* for 15 min at 4 °C. The resulting mouse serum supernatant was analyzed via Sepp1 (Elabscience, Cat. E-EL-M3044) and S100a9 (Elabscience, Cat. E-EL-M3049) ELISA kits according to the manufacturer’s instructions. SAM concentrations were determined using the ELISA Kit for S-Adenosyl Methionine (Cloud-Clone Corp, Cat. CEG414Ge) according to the manufacturer’s instructions. For human samples, whole blood was collected from healthy individuals and HCC patients following ethical approval and informed consent. These samples were processed to sera, which were aliquoted and stored at −80 °C pending analysis. The human serum samples were analyzed using SEPP1 (Elabscience, Cat. E-EL-H2177) and S100A9 (Elabscience, Cat. E-EL-H6197) ELISA kits according to the manufacturer’s instructions. For assessments of spent medium samples, ELISA was performed using IFNγ (Multi Sciences, Cat. EK280) and IL2 (Multi Sciences, Cat. EK202) ELISA kits according to the manufacturer’s instructions.

### Analysis of serum AST, ALT, and GPx levels

Serum levels of aspartate aminotransferase (AST) and alanine aminotransferase (ALT) (Shanghai Shensuo, 3040280 and 3050280), as well as that of glutathione peroxidase (GPx) (Nanjing Jiancheng Bioengineering Institute, A005-1-2), were measured using commercially available assay kits according to the respective manufacturer’s instructions.

### Western blotting

Protein from cells or tissues was extracted using radioimmunoprecipitation assay (RIPA) buffer (150 mM NaCl, 20 mM Tris-HCl pH7.4, 1 mM ethylenediaminetetraacetic acid (EDTA), 1% Triton-X100, 0.1% sodium dodecyl sulfide (SDS) containing of protease and phosphatase inhibitors). Prior to loading onto gels, samples were denatured by boiling at 100 °C for 10 min in 5× SDS protein sample loading buffer. Volumes of lysates containing equivalent amounts of total protein then were subjected to SDS-polyacrylamide gel electrophoresis (PAGE) and transferred to polyvinylidene fluoride (PVDF) membranes, which then were blocked with 5% non-fat milk for 1 h at room temperature. The blocked membranes were probed with the indicated primary antibodies, followed by the appropriate horse radish peroxidase (HRP) -conjugated secondary antibodies.

The primary antibodies used in the experiments were antibodies with specificity for SEPP1 (1:300, Santa Cruz, Cat. sc-376858), histone H3 (1:1000, Cell Signaling Technology, Cat. 9715S), acetyl-histone H3 (Lys27) (1:1000, Cell Signaling Technology, Cat. 8173S), histone H3 (trimethyl Lys4) (1:1000, Novus Biologicals, Cat. NB21-1023), histone H3 (trimethyl Lys27) (1:1000, Selleck, F0165), p21 (1:1000, Abcam, Cat. ab188224), HSP90 (1:1000, Cell Signaling Technology, Cat. 4877S), and GAPDH (1:1000, Proteintech, Cat. 1E6D9; used as the loading control). Uncropped images of all immunoblots are provided in the Supplementary Fig.

### Immunofluorescence staining

Mouse tumor tissues were fixed in 4% paraformaldehyde for 24 h, then trimmed, embedded in paraffin, sectioned, and placed on slides. Sections were deparaffinized and rehydrated in xylene and ethanol. Antigen retrieval was performed in Tris-EDTA Buffer (10 mM Tris, 1 mM EDTA, pH 9.0) for 15 min in the microwave. Tumor sections were blocked with 1% bovine serum albumin (BSA) for 30 min at room temperature, after which sections were incubated (overnight at 4 °C) in PBS-1% BSA containing a primary antibody against green fluorescent protein (GFP; 1:500, Abmart, Cat. M20004), Ly6G (1:200, Servicebio, Cat. GB11229), or S100A9 (1:500, R&D Systems, Cat. AF2065). Sections then were stained (for 2 h at room temperature, in the dark) in PBS-1% BSA containing a secondary antibody consisting of (as appropriate) donkey anti-mouse IgG (H+L) Alexa Fluor 488 (1:500, Thermo Fisher Scientific, Cat. A-21202), donkey anti-goat IgG (H+L) Alexa Fluor 568 (1:500, Thermo Fisher Scientific, Cat. A-11057), or chicken anti-rabbit IgG (H+L) Alexa Fluor 647(1:500, Thermo Fisher Scientific, Cat. A-21443). Finally, the sections were counterstained for cell nuclei (DNA) with 4′,6-diamidino-2-phenylindole (DAPI; 5 μg/mL, Thermo Fisher, Cat. 62248) and mounted using Fluoromount™ Aqueous Mounting Medium (Sigma Aldrich, Cat. F4680). Fluorescent images were obtained with an Olympus FV1000 confocal laser-scanning microscope.

### CUT&RUN assay for chromatin profiling

CUT&RUN was performed to map H3K4me3-marked regions of the genome with high resolution and low background. Briefly, TINs were isolated using magnetic beads, and digitonin was used to permeabilize the cells. Permeabilized cells then were incubated with a primary antibody specific to H3K4me3 (Novus Biologicals, Cat. NB21-1023), H3K27ac (CST, Cat. 4353S) and H3K9me2 (Novus Biologicals, Cat. NB21-1072) followed by incubation with protein A-micrococcal nuclease (pA-MNase; Beyotime, Cat. D7195). Upon activation with calcium, MNase selectively cleaves accessible DNA proximal to the H3K4me3-marked nucleosomes. Released DNA fragments were extracted and prepared for sequencing using an Illumina platform. Libraries were sequenced in paired-end mode (PE150), generating a total of ∼ 3 Gb of data per sample.

After sequencing, raw reads were processed for quality control using fastp, and the cleaned reads were aligned to the reference genome using Bowtie2 with default parameters optimized for CUT&RUN data. Aligned reads were filtered to remove duplicates and low-quality alignments using SAMtools. To identify and mark duplicate sequences in the aligned data (given that duplicate reads might otherwise create potential biases), we employed GATK’s Mark Duplicates. Peaks representing H3K4me3-enriched regions were identified using MACS2 in broad peak-calling mode. Genome-wide visualization was achieved by generating normalized bigWig files using deepTools; the resulting files were loaded into Integrative Genomics Viewer (IGV) for interactive exploration. Additionally, heatmaps and metagene plots were generated using deepTools to provide a global overview of H3K4me3 distribution relative to genomic features.

### Single-cell RNA sequencing

scRNA-seq was performed using the 10x Genomics Chromium platform with the 3’ gene expression kit (v. 3). Briefly, fresh tumor tissues were excised from mouse liver cancer models and enzymatically dissociated. The resulting single-cell suspensions were filtered through a 40-µm cell strainer, and cell viability was assessed to ensure values of >90% before proceeding. Approximately 10,000 cells per sample were captured and processed for library construction according to the manufacturer’s protocol. Sequencing was conducted on an Illumina platform with a target depth of 50,000 reads per cell. Raw data were processed using Cell Ranger (v7.1.0) for alignment to the mouse genome and for generation of the gene-cell matrix. Downstream analysis, including quality control, normalization, clustering, and cell type annotation, was performed using Seurat (v. 4.0); cells showing high mitochondrial gene expression or low Unique Molecular Identifier (UMI) counts were excluded.

### Survival and expression analysis

The effects of SEPP1 and S100A9 on the survival of patients with breast cancer were analyzed using the Kaplan–Meier Plotter (http://kmplot.com/analysis/) online tool. The levels of SEPP1 protein in tumors and adjacent normal tissues were analyzed using data from the National Cancer Institute’s CPTAC database, as accessed (on April 9, 2025) via The University of Alabama at Birmingham Cancer Data Analysis Portal (UALCAN) (http://ualcan.path.uab.edu/). Gene expression correlation of *SEPP1* and *S100A9* analysis was performed using the TIMER database (http://timer.cistrome.org/).

### Human samples

Human tissues and adjacent normal tissues were collected from Changhai Hospital, with patient consent. Following collection, specimens were snap-frozen in liquid nitrogen and store at - 80 ℃ pending analysis. Blood was collected from Changhai Hospital with patient consent. The use of all clinical tissue samples in this study was approved by the Ethics Committees at the Shanghai Institute of Nutrition and Health, Chinese Academy of Sciences and the Changhai Hospital (Approval No. CHEC2024-180).

### Statistical analyses

Data analyses were performed using Prism 8.0 (GraphPad Software, La Jolla, CA, USA). The data presentation and statistical analyses are described in the figure legends. Where appropriate, analyses were performed as two-tailed tests. p <0.05 was considered statistically significant.

## Acknowledgment

This study was supported by the National Key Research and Development Program of China (2021YFA1100504 to L.Z.) and Program of EnShi TuJia & Miao Autonomous Prefecture Bureau of Scientific & Technological Affair (L.Z.)

## Author contributions

L.Z. supervised this work. L.Z. and J.J. designed the research and wrote, reviewed, and edited the manuscript. J.J. performed most experiments. K.X., R.C. and KY.X. performed a specific subset of the experiments and analyses. X.X. provided constructive suggestions for the design of the research.

## Competing interests

The authors declare no potential conflicts of interest.

---

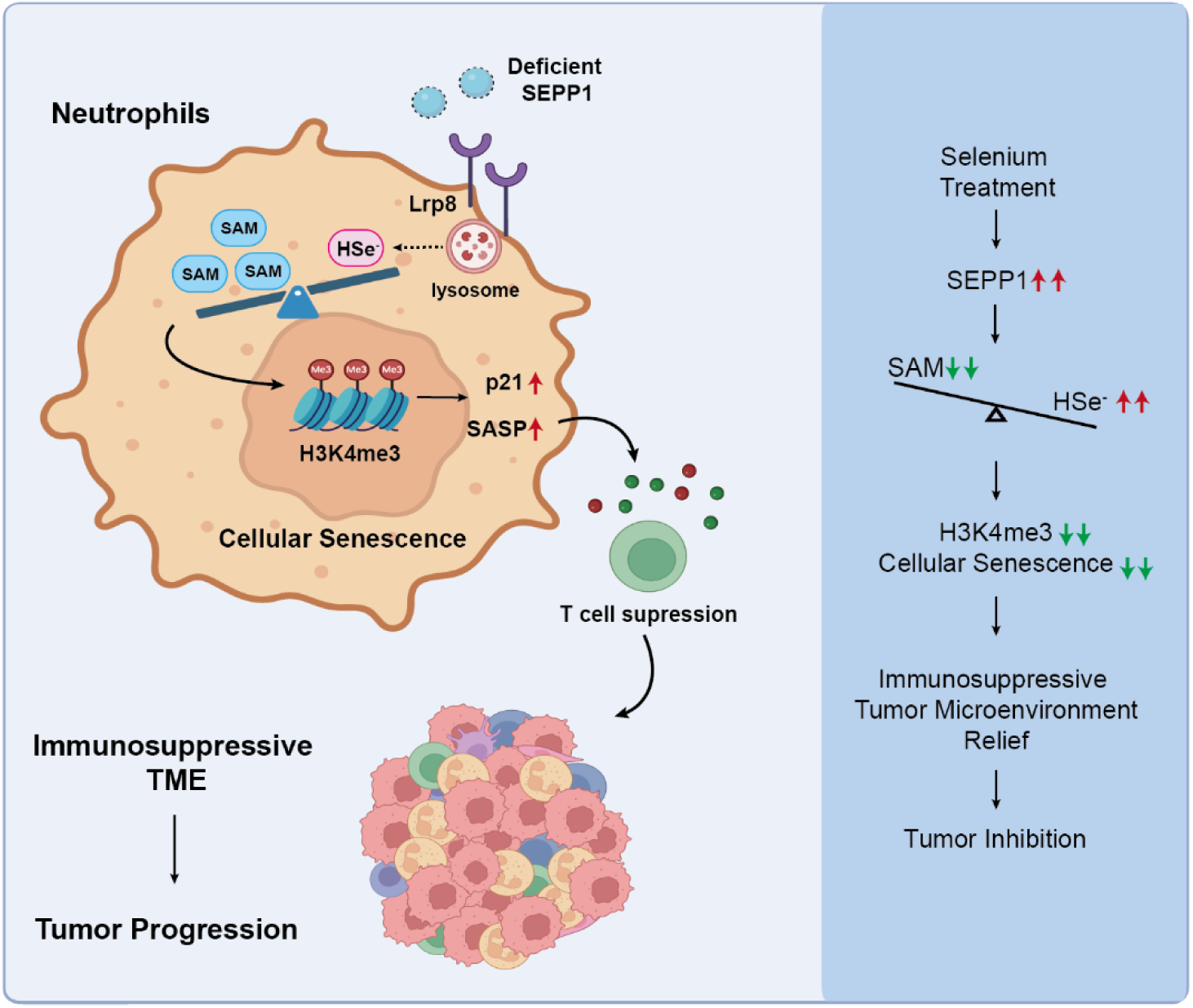
Working Model.

**Supplementary Figure 1:**
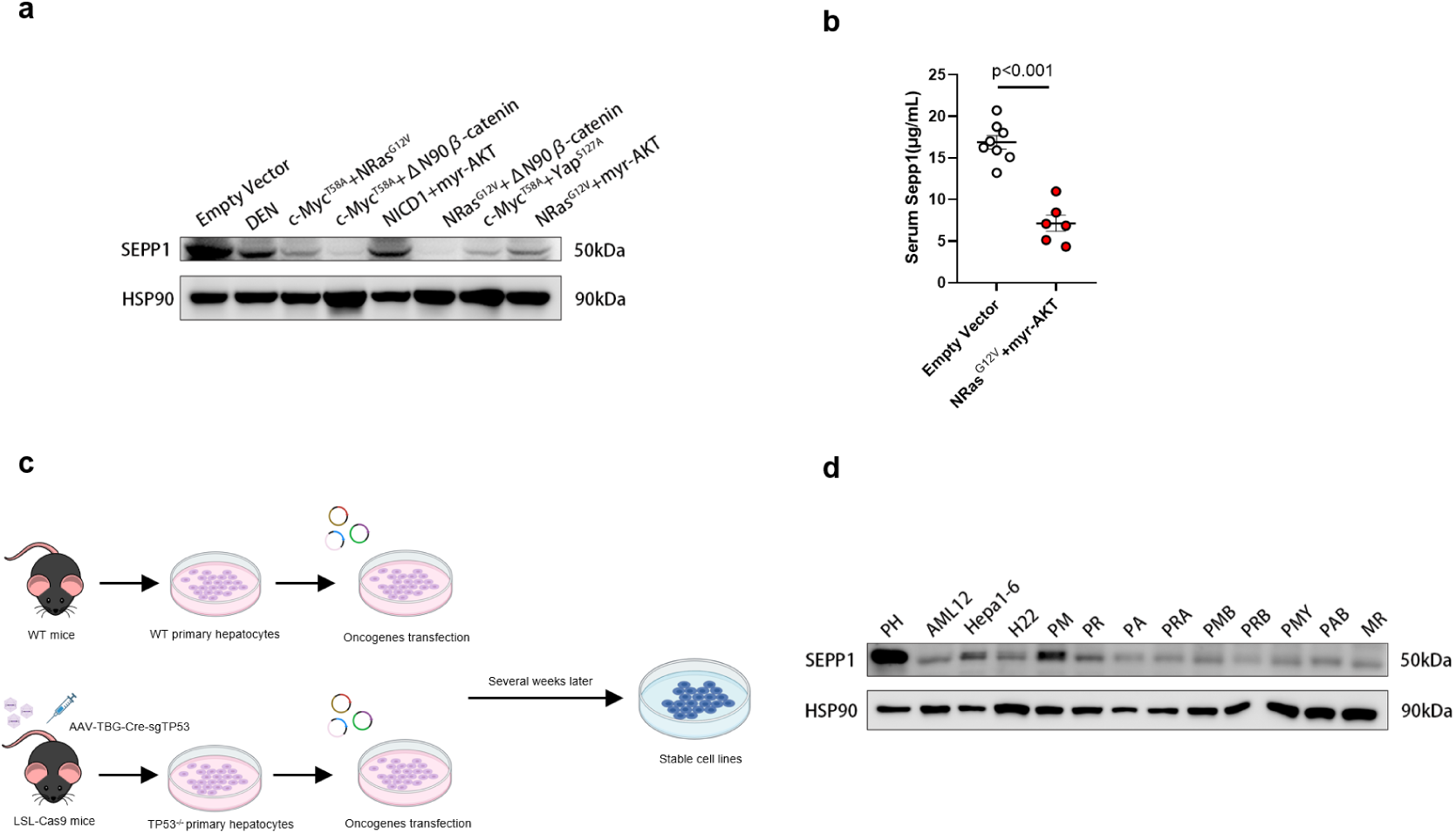
Downregulation of Sepp1 in various mouse liver cancer contexts. (a) Western blot analysis of Sepp1 protein levels in tumor tissues from mice injected with different oncogenic constructs. (b) Serum Sepp1 protein levels, as measured by ELISA, in control mice (injected with empty vector) and in a liver cancer mouse model (transfected with constructs encoding NRas^G12V^ and myr-AKT proteins) (n = 6–8 per group). (c) Schematic representation of the generation of stable liver cancer cell lines. To facilitate further analysis, stable cell lines were established by transfecting primary hepatocytes from wild-type (WT) or *p53 tumor suppressor* (*Trp53*) knockout mice with constructs encoding oncogenic proteins. (d) Western blot analysis of Sepp1 protein in primary hepatocytes and multiple stable liver cancer cell lines established with different oncogenes. Data are presented as mean ± SEM; statistical significance was determined using two-tailed unpaired Student’s t-test.

**Supplementary Figure 2:**
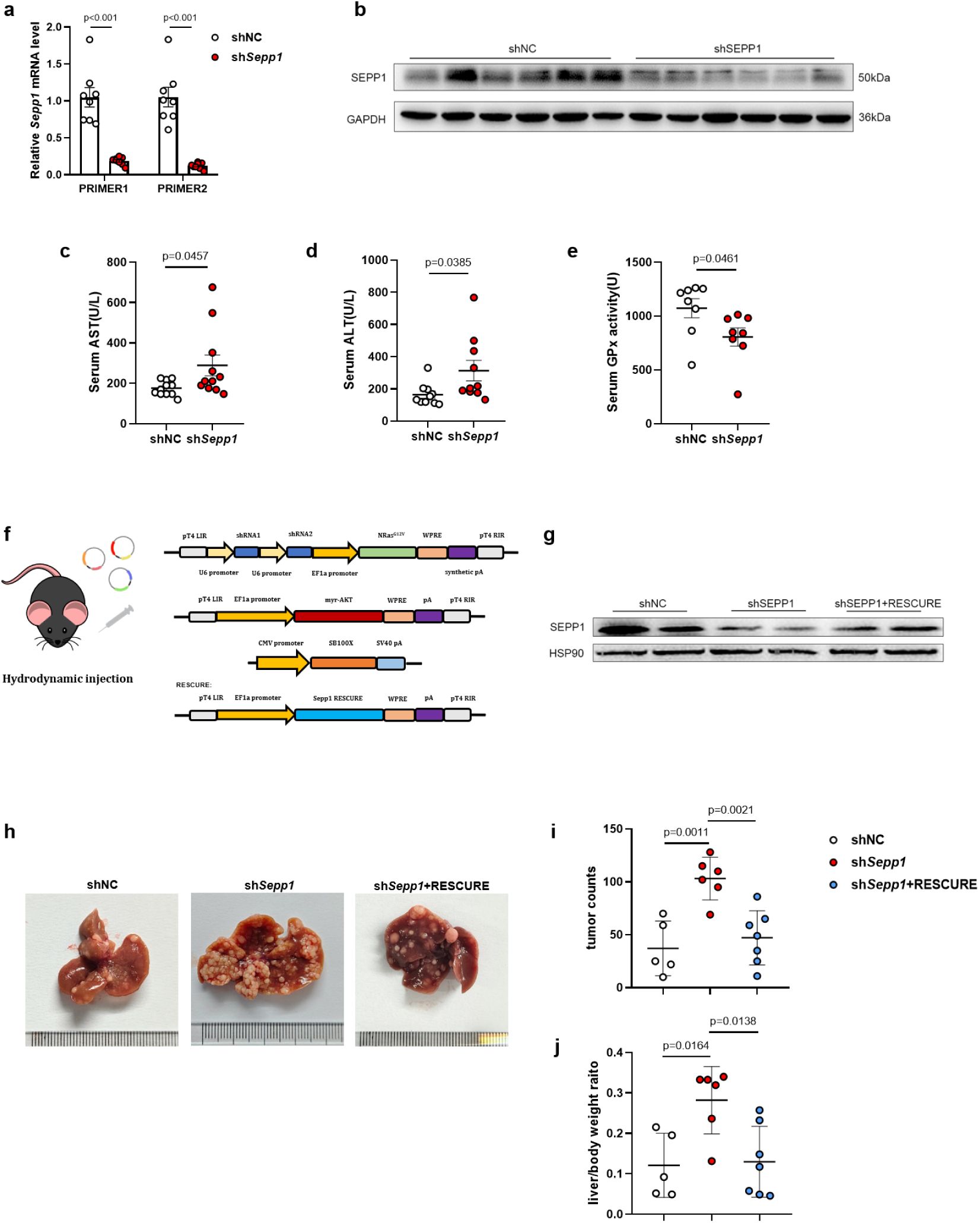
Validation and rescue experiment to assess the impact of *Sepp1* knockdown on tumor burden. (a) Quantitative RT-PCR analysis of *Sepp1* mRNA levels using two different primer sets (PRIMER1 and PRIMER2) in shNC and sh*Sepp1* liver tumor tissues, confirming significant knockdown of *Sepp1* in sh*Sepp1* mice compared to shNC controls (n = 8 per group). (b) Western blot analysis of Sepp1 protein levels in shNC and sh*Sepp1* liver tumor tissues; Glyceraldehyde 3-phosphate dehydrogenase (GAPDH) was used as a loading control. (c–e) Serum biochemical analysis from tumor-bearing mice to assess liver function. (c) Serum aspartate aminotransferase (AST) levels, (d) Serum alanine aminotransferase (ALT) levels, and (e) Serum glutathione peroxidase (GPx) activity levels were measured, showing significant increases in AST and ALT, and a significant decrease in GPx activity, in sh*Sepp1* mice compared to shNC animals (n = 8–11 per group, p-values as indicated). (f) Schematic of hydrodynamic injection constructs used in the rescue experiment. Mice were injected with construct encoding oncogenic NRas^G12V^ and myr-AKT protein along with either shNC, sh*Sepp1*, or sh*Sepp1* plus a *Sepp1* rescue construct (*Sepp1*-RESCUE). (g) Representative western blot analysis showing Sepp1 protein levels in shNC, sh*Sepp1*, and sh*Sepp1*+RESCUE groups. (h) Representative images of liver morphology from the respective groups: control (shNC), *Sepp1* knockdown (sh*Sepp1*), and *Sepp1* knockdown with *Sepp1* rescue (sh*Sepp1*+RESCUE). *Sepp1* knockdown resulted in increased tumor burden, which was alleviated by *Sepp1* rescue. (i) Tumor counts in liver tissues from shNC, sh*Sepp1*, and *Sepp1*-RESCUE groups. sh*Sepp1* mice exhibited significantly increased tumor counts compared to shNC controls, while *Sepp1*-RESCUE mice showed decreased tumor counts compared to sh*Sepp1* mice (n = 6–7 per group). (j) Liver-to-body weight ratio in shNC, sh*Sepp1*, and *Sepp1*-RESCUE mice (n = 6–7 per group). Data are presented as mean ± SEM; statistical significance was determined using two-tailed unpaired Student’s t-tests (a, c-e) or two-tailed One-way ANOVA with post hoc Tukey multiple comparisons tests (i, j).

**Supplementary Figure 3:**
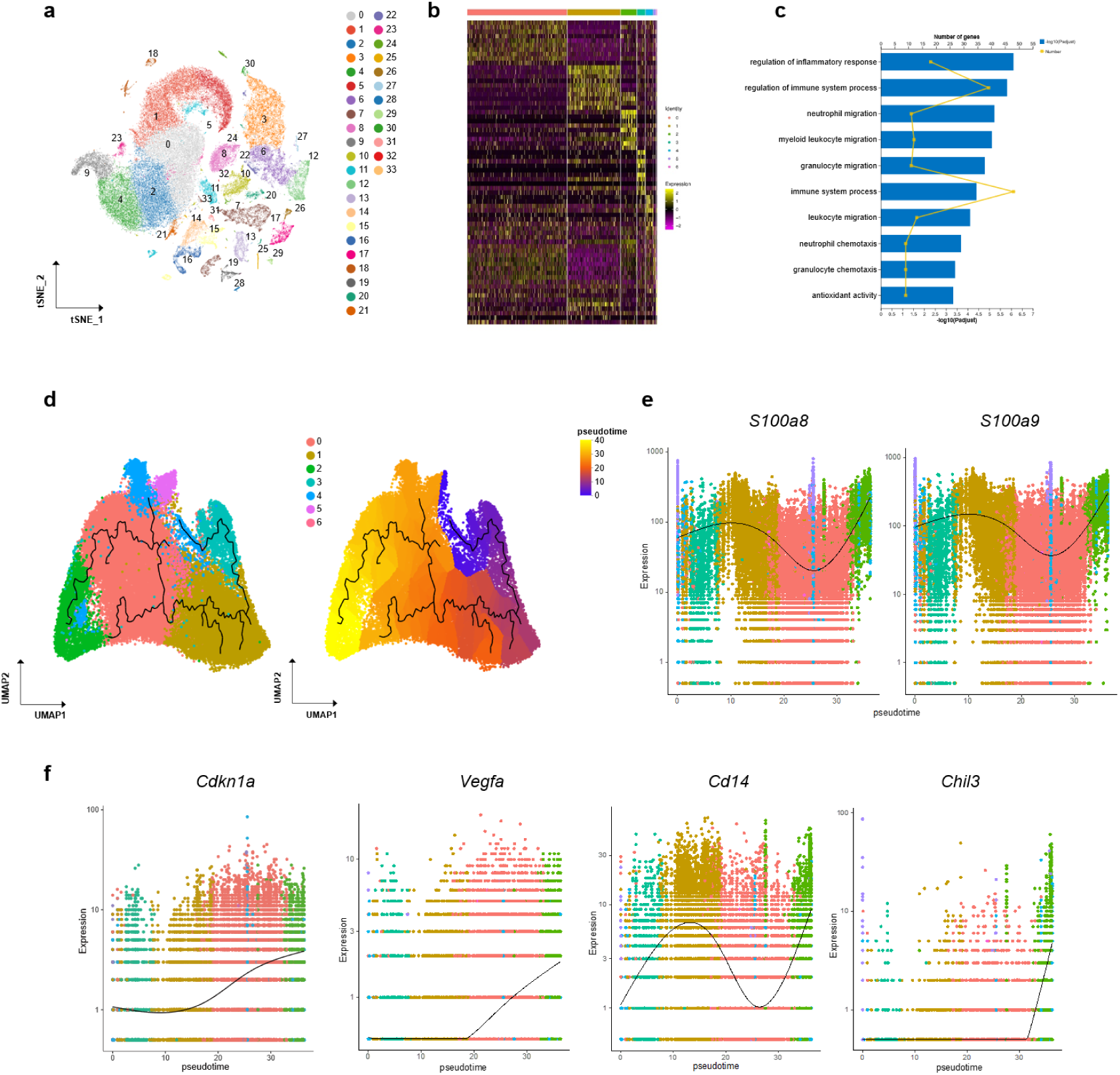
scRNA-seq analysis of tumor-associated neutrophils in *Sepp1* knockdown and control conditions. (a) t-SNE plot of single-cell RNA sequencing (scRNA-seq) data from liver tumors. (b) Heatmap showing differentially expressed genes across 7 identified neutrophil subclusters identified via scRNA-seq analysis. (c) Gene enrichment analysis of genes upregulated in Subcluster 2. (d) Pseudotime analysis (using Monocle3 software) of all neutrophils revealed that Subcluster 2 belongs to the latest stage in the pseudotime trajectory, indicating that this subcluster differentiates from other subclusters, an inference that is consistent with the senescent characteristics of Subcluster 2. (e-f) Kinetics plot illustrating the expression levels of *S100a8/9*, *Cdkn1a*, *Vegfa*, *Cd14*, and *Chil3* across the pseudotime-course of neutrophil subclusters. The black smooth line depicts the trend of gene expression as this parameter varies along pseudotime.

**Supplementary Figure 4:**
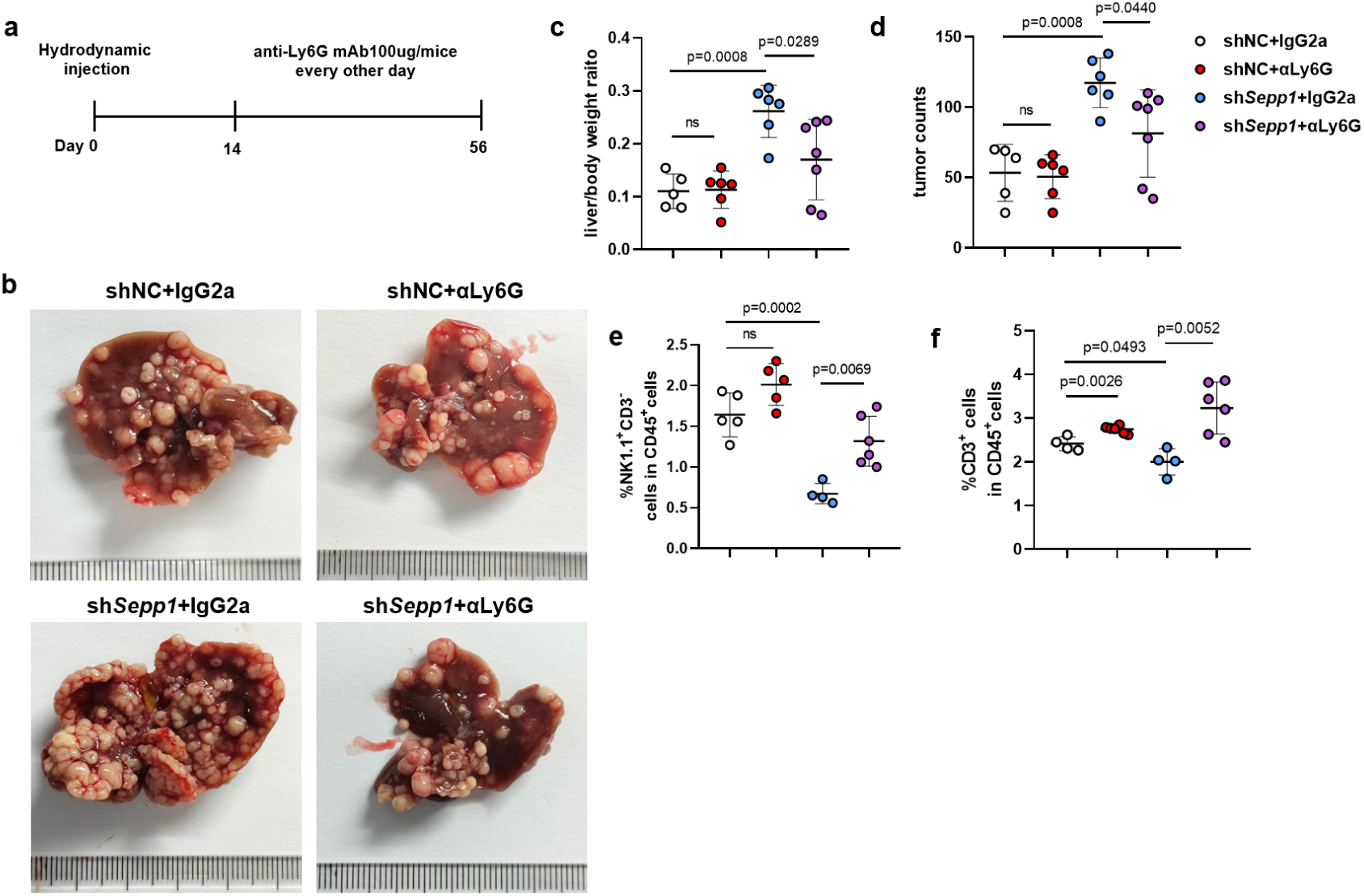
Neutrophil depletion mitigates tumor burden in Sepp1 knockdown liver tumors. (a) Experimental timeline for the treatment with antibody against lymphocyte antigen 6 family member G (Ly6G). Mice injected via hydrodynamic tail vein (HDTVi) with constructs encoding oncogenic NRas^G12V^ and myr-AKT protein, along with shNC or sh*Sepp1* constructs, were treated (from Day 14 to Day 56) with anti-Ly6G monoclonal antibody (mAb) (100 µg/mouse every other day) to deplete neutrophils. (b) Representative images of livers from different treatment groups. (c, d) Quantification of liver-to-body weight ratio (c) and tumor counts (d) for each group. Neutrophil depletion significantly decreased relative liver weight and tumor numbers in sh*Sepp1* mice treated with anti-Ly6G antibody (αLy6G) compared to the isotype (immunoglobulin 2a; IgG2a) control (n = 5–7 per group). (e) Flow cytometric analysis showing the percentage of natural killer (NK) cells in CD45^+^ cells. Neutrophil depletion significantly increased the proportion of NK cells in sh*Sepp1* groups (n = 4-6 per group). (f) Flow cytometric analysis showing the percentage of T cells in CD45^+^cells. Depletion of neutrophils significantly increased the infiltration of T cells (n = 4-6 per group). Data are presented as mean ± SEM; statistical significance was determined using two-tailed One-way ANOVA with post hoc Tukey multiple comparisons tests (c, d, e) or two-tailed unpaired Student’s t-test (f).

**Supplementary Figure 5:**
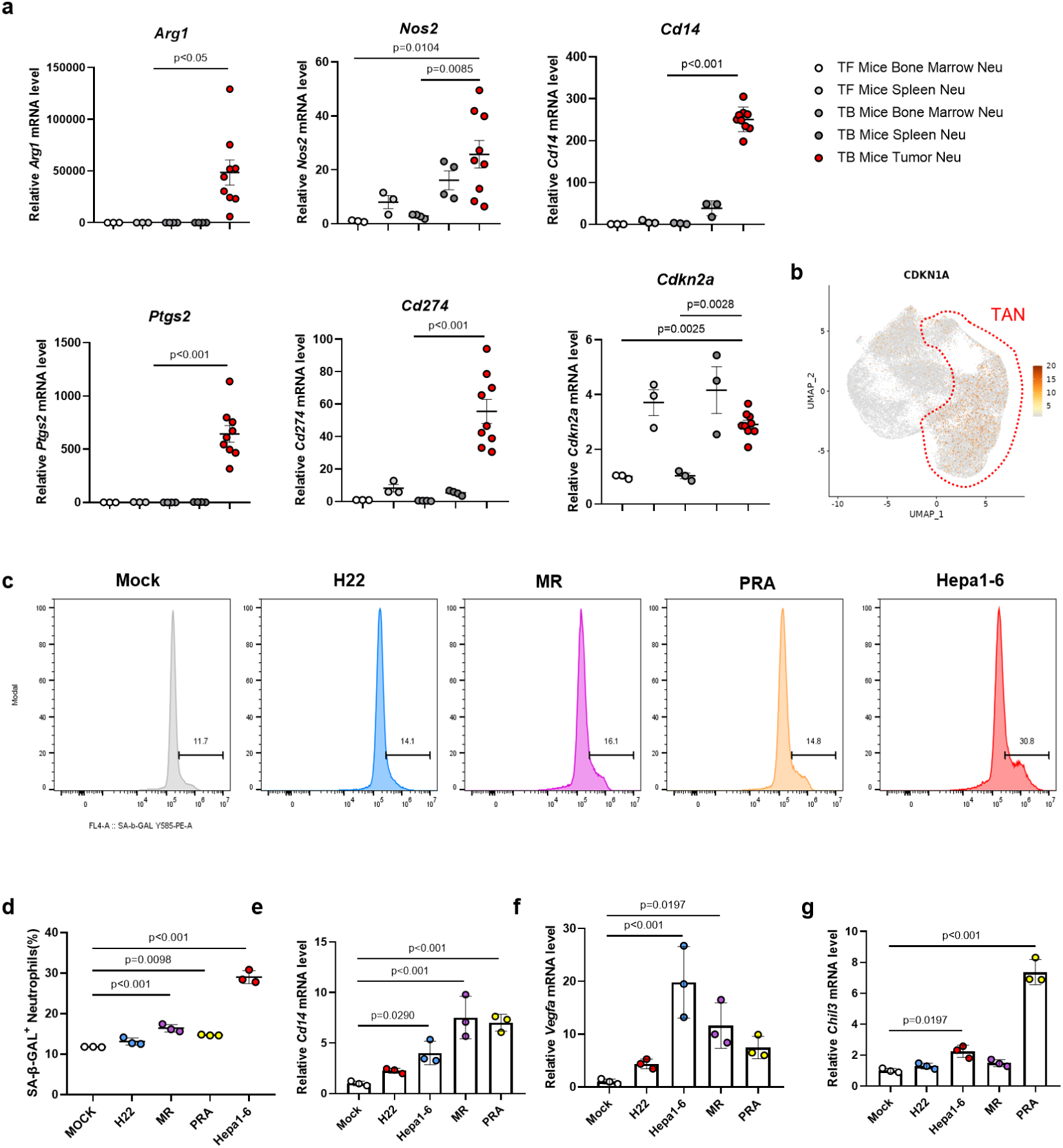
p21^+^ senescent-like neutrophils are present exclusively in tumor tissues. (a) Quantitative RT-PCR analysis of gene expression in neutrophils isolated from different origins in tumor-free (TF) and tumor-bearing (TB) mice, including bone marrow (BM), spleen, and tumor tissues. The transcripts of well-known immunosuppression genes (including *Arg1*, *Nos2*, *Cd14*, *Ptgs2*, and *Cd274*) accumulated to significantly higher levels in tumor-associated neutrophils (TANs; TB Mice Tumor Neu) compared to neutrophils from other sources. Similarly, *Cdkn2a* mRNA accumulated to significantly higher levels in tumor neutrophils compared to those from the BM of TF or TB mice (n = 3–9 per group). (b) Uniform Manifold Approximation and Projection (UMAP) plot highlighting the expression of *Cdkn1a* in TANs. The region marked as “TAN” (red dashed line) indicates a subset of neutrophils present in tumors but not in peripheral blood or peritumoral tissues. (Data from http://meta-cancer.cn:3838/scPLC/.) (c) Flow cytometric analysis showing the percentage of senescence-associated beta-galactosidase (SA-β-GAL) -positive neutrophils following treatment with conditioned medium from various liver cancer cell lines, including mock (control), H22, MR, PRA, and Hepa1-6. (d) Quantification of SA-β-GAL-positive neutrophils from flow cytometric data, indicating that conditioned medium from MR, PRA, and Hepa1-6 cell lines significantly induces senescence in primary neutrophils compared to the mock control (n = 3 per group). Data are presented as mean ± SEM; statistical significance was determined using the Student’s t-test. (e-g) Quantitative RT-PCR analysis of transcript levels in primary neutrophils treated with conditioned medium from liver cancer cell lines. *Cd14*, *Vegfa*, and *Chil3* mRNA levels were significantly elevated in neutrophils exposed to conditioned medium with the H22, MR, PRA, and Hepa1-6 cell lines compared to the mock control (n = 3 per group). Data are presented as mean ± SEM; statistical significance was determined using One-way ANOVA with Dunnett multiple comparisons tests.

**Supplementary Figure 6:**
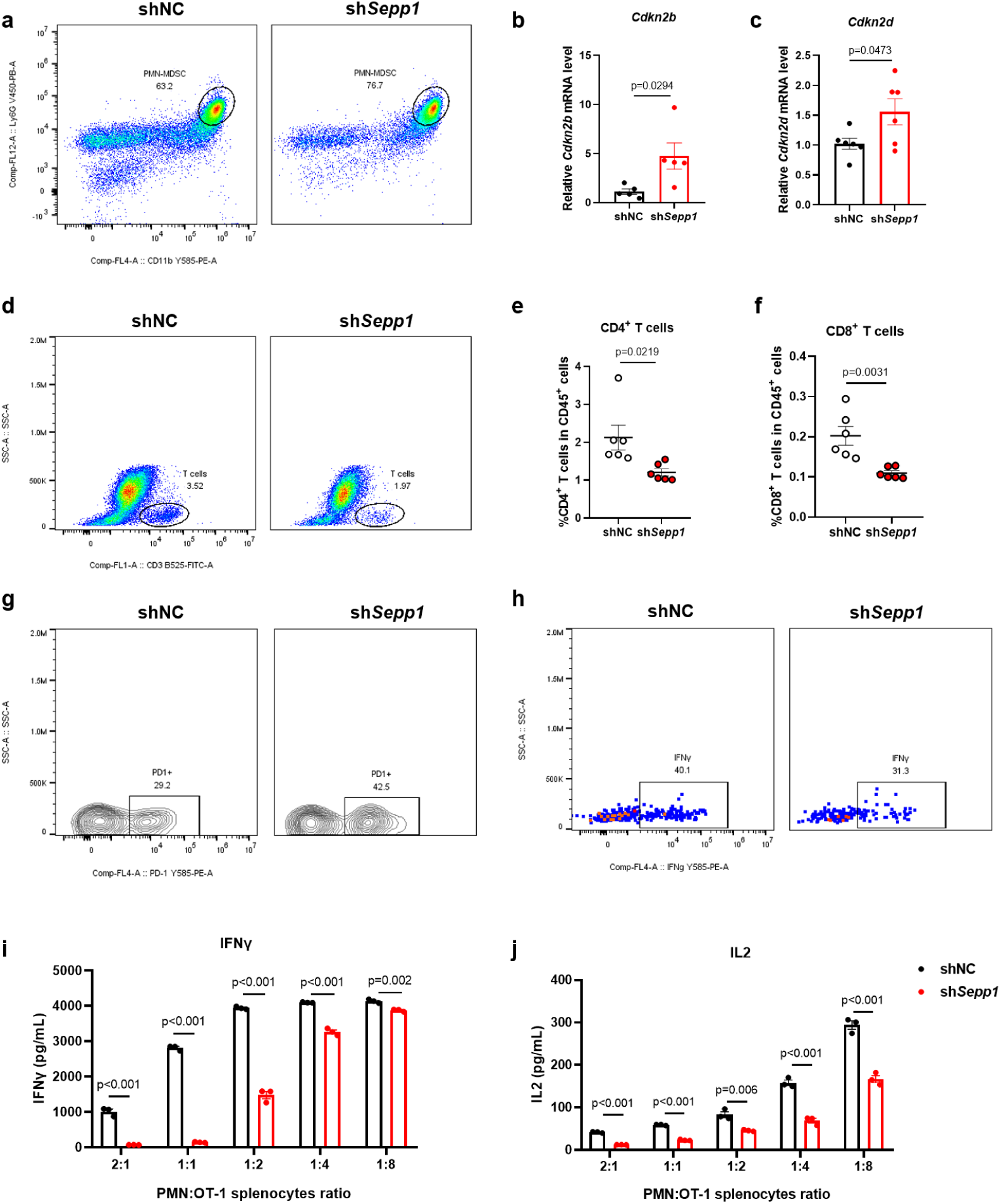
*Sepp1* deficiency induces neutrophil senescence and immunosuppression, impeding T cell infiltration and activity in tumors (related to Figure 2). (a) Flow cytometric analysis of polymorphonuclear myeloid-derived suppressor cells (PMN-MDSCs) from shNC and sh*Sepp1* mice. (b, c) Relative mRNA levels of *Cdkn2b* (b) and *Cdkn2d* (c) in neutrophils isolated from shNC and sh*Sepp1* animals (n=5-6 per groups). (d) Flow cytometric analysis of T cell proportions in shNC and sh*Sepp1* mice. (e-f) CD4^+^ T cells and CD8^+^ T cells proportions in the tumor microenvironment of shNC and sh*Sepp1* mice. (g) Programmed cell death protein 1 (PD-1) expression in CD8^+^ T cells, as assessed by flow cytometry. (h) Cytokine staining of intracellular interferon gamma (IFNγ) in CD8^+^ T cells from shNC and sh*Sepp1* mice. (i, j) ELISA quantification of IFNγ (d) and IL-2 (e) in the co-culture spent medium, demonstrating significantly decreased levels of both of these cytokines in cultures employing neutrophils isolated from *Sepp1*-depeleted tumors (compared to neutrophils isolated from control tumors) across various neutrophils: OT-1 splenocyte ratios (n = 3 per group). Data are presented as mean ± SEM; statistical significance was determined using two-tailed unpaired Student’s t-tests.

**Supplementary Figure 7.**
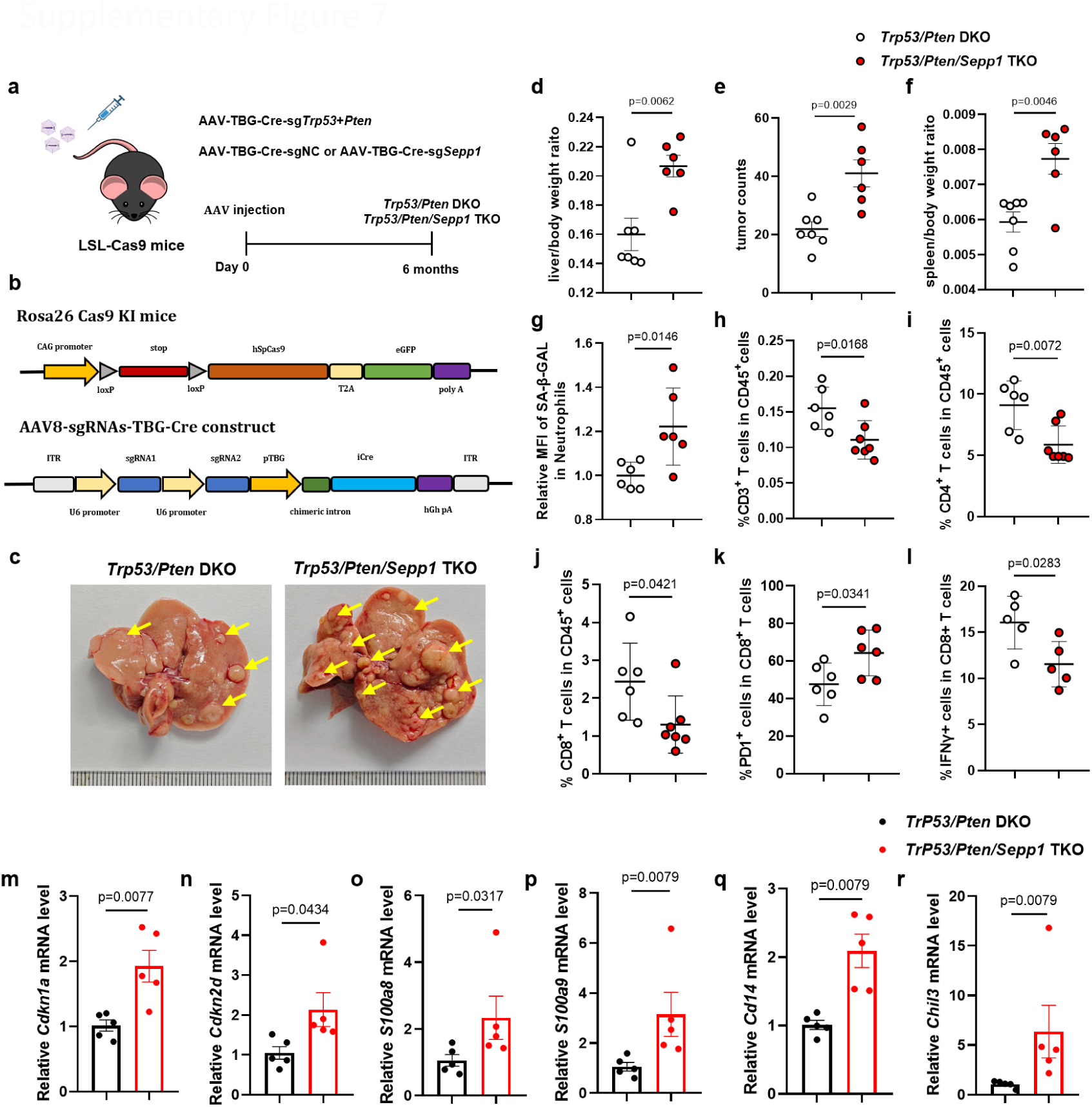
*Sepp1* knockout significantly potentiates tumor burden in *Trp53*/*Pten* double-knockout mice with hepatocellular carcinoma. (a) Schematic representation of the experimental design. LSL-Cas9 (*Cas9* knock-in mice; genotypically *Rosa26^Cas9+/+^*) knock-in mice were injected with AAV-TBG-Cre-sg*Trp53*+*Pten* and AAV-TBG-Cre-sgNC to generate *p53 tumor suppressor protein* (*Trp53) / phosphatase and tensin homolog* (*Pten*) double-knockout (DKO) mice, or with AAV-TBG-Cre-sg*Trp53*+*Pten* and AAV-TBG-Cre-sg*Sepp1* to generate *Trp53/Pten/Sepp1* triple-knockout (TKO) mice. Tumor burden and immune responses were analyzed 6 months post-injection. (b) Schematic diagrams of the gene order and constructs of *Rosa26^Cas9+/+^*knock-in (KI) mice and the AAV8-sgRNA-TBG-Cre system used for gene knockout. (c) Representative liver images from *Trp53/Pten* DKO and *Trp53/Pten/Sepp1* TKO mice. Yellow arrows indicate tumor nodules, the number of which were significantly increased in TKO mice (compared to DKO mice). (d-f) Quantification of liver-to-body weight ratios (d), tumor counts (e), and spleen-to-body weight ratios (f), showing significantly increased relative liver weight, tumor burden, and relative spleen weight in TKO mice compared to DKO mice (n = 6-7 per group). (g) Flow cytometric analysis of senescence-associated beta-galactosidase (SA-β-GAL) mean fluorescence intensity (MFI) in neutrophils from the various mice, indicating increased senescence in neutrophils from TKO mice compared to those from DKO mice. (h) Flow cytometric quantification of CD3^+^ T cells in the tumor microenvironment of DKO and TKO mice, indicating significantly decreased T cell infiltration in tumors from TKO mice compared to those from DKO mice. (i-j) Proportions of CD4⁺ and CD8⁺ T cells in CD45⁺ cells; values were significantly decreased in cells from TKO mice compared to those from DKO mice (k-l) Immune functional analysis showing an increase in PD-1⁺ exhausted CD8⁺ T cells and a decrease in IFN-γ⁺ CD8⁺ T cells, suggesting impaired anti-tumor immunity in TKO mice compared to DKO mice. (m-r) RT-qPCR analysis of the levels of transcripts of inflammatory and immune-related genes (*Cdkn1a*, *Cdkn2d*, *S100a8*, *S100a9*, *Cd14*, *Chil3*) in magnetic-activated cell sorting (MACS). neutrophils, showing significantly increased levels of these transcripts in TKO mice compared to DKO mice (n = 5 per group). Data are presented as mean ± SEM, with statistical significance determined by two-tailed unpaired Student’s t-tests or Mann-Whitney U-tests.

**Supplementary Figure 8:**
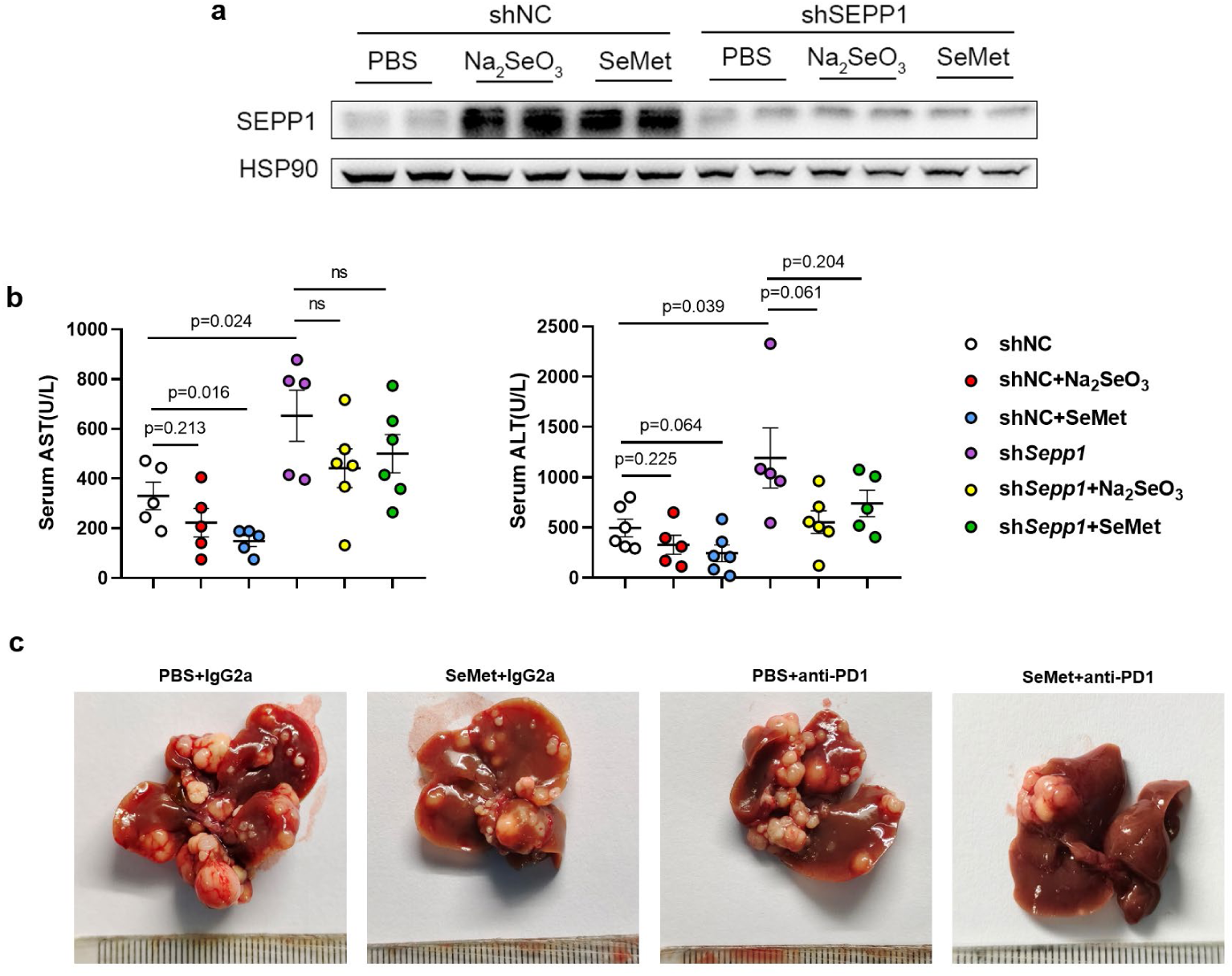
Selenium inhibits tumor growth in a Sepp1-dependent manner (related to Figure 4). (a) Selenium supplementation significantly increased the protein levels of Sepp1 in tumors from shNC mice (compared to those seen in the absence of Se supplementation); this potentiation was not observed in sh*Sepp1* mice. (b) Serum aspartyl transferase (AST) and alanyl transferase (ALT) levels in the various treatment groups. (c) Representative images of liver tumors from selected treatment groups, including animals receiving phosphate-buffered saline (vehicle) + isotype antibody (immunoglobulin G2a; IgG2a); selenomethionine (SeMet) + IgG2a; PBS + anti-programmed cell death protein 1 antibody (anti-PD-1); or SeMet + anti-PD-1. Images show the macroscopic appearance of tumor burden under the various conditions. Data are presented as mean ± SEM; statistical significance was determined using two-tailed unpaired Student’s t-test.

**Supplementary Figure 9:**
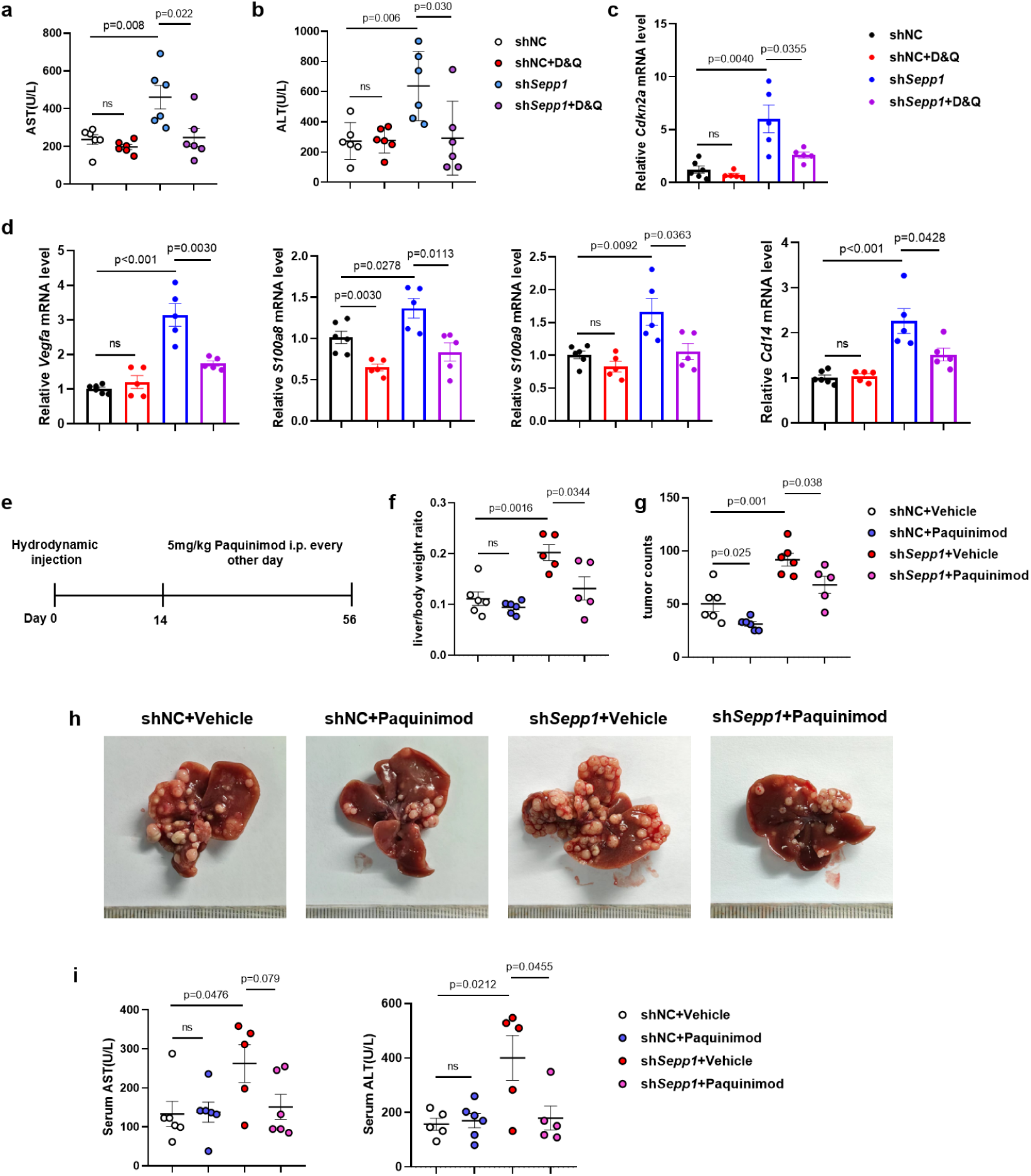
The effects of D&Q and paquinimod treatments on tumor burden and associated markers in liver tumors from *Sepp1* knockdown and control mice. (a-b) Serum levels of aspartyl transferase (AST) (a) and alanyl transferase (ALT) (b), showing the circulating levels of these liver injury markers across treatment groups. Treatment with dasatinib and quercetin (D&Q) provided significantly decreased AST and ALT levels in the sh*Sepp1* mice (compared to levels in sh*Sepp1* animals dosed with vehicle); this effect was not seen in shNC mice. (c–d) Relative mRNA levels of *Cdkn2a* (c), *Vegfa*, *S100a8*, *S100a9*, and *Cd14* (d) in tumors, as measured by RT-qPCR. For each of these markers, D&Q treatment provided significantly decreased transcript levels in the sh*Sepp1* mice (compared to levels in sh*Sepp1* animals dosed with vehicle); among the shNC mice, this effect was seen only for *S100a8* transcript levels. (e) Experimental schematic timeline for paquinimod treatment. Mice were subjected to hydrodynamic tail vein injection (HDTVi) with constructs encoding oncogenic NRas^G12V^ and myr-AKT protein in combination with shNC or sh*Sepp1* constructs; animals subsequently were treated (Day 14 to Day 56 post-HDTVi) with paquinimod (5 mg/kg, administered intraperitoneally every other day). (f, g) Liver-to-body weight ratios (f) and tumor counts (g), indicating that (compared to vehicle) treatment with paquinimod significantly attenuated relative liver weight in sh*Sepp1* mice (but not in shNC mice), and tumor burden in both sh*Sepp1* and shNC mice (n = 6–7 per group, p-values as indicated). (h) Representative images of livers from shNC+Vehicle, shNC+paquinimod, sh*Sepp1*+Vehicle, and sh*Sepp1*+paquinimod groups. (i) Serum levels of AST and ALT in paquinimod treatment groups. Data are presented as mean ± SEM; statistical significance was determined using two-tailed unpaired Student’s t-tests.

**Supplementary Figure 10:**
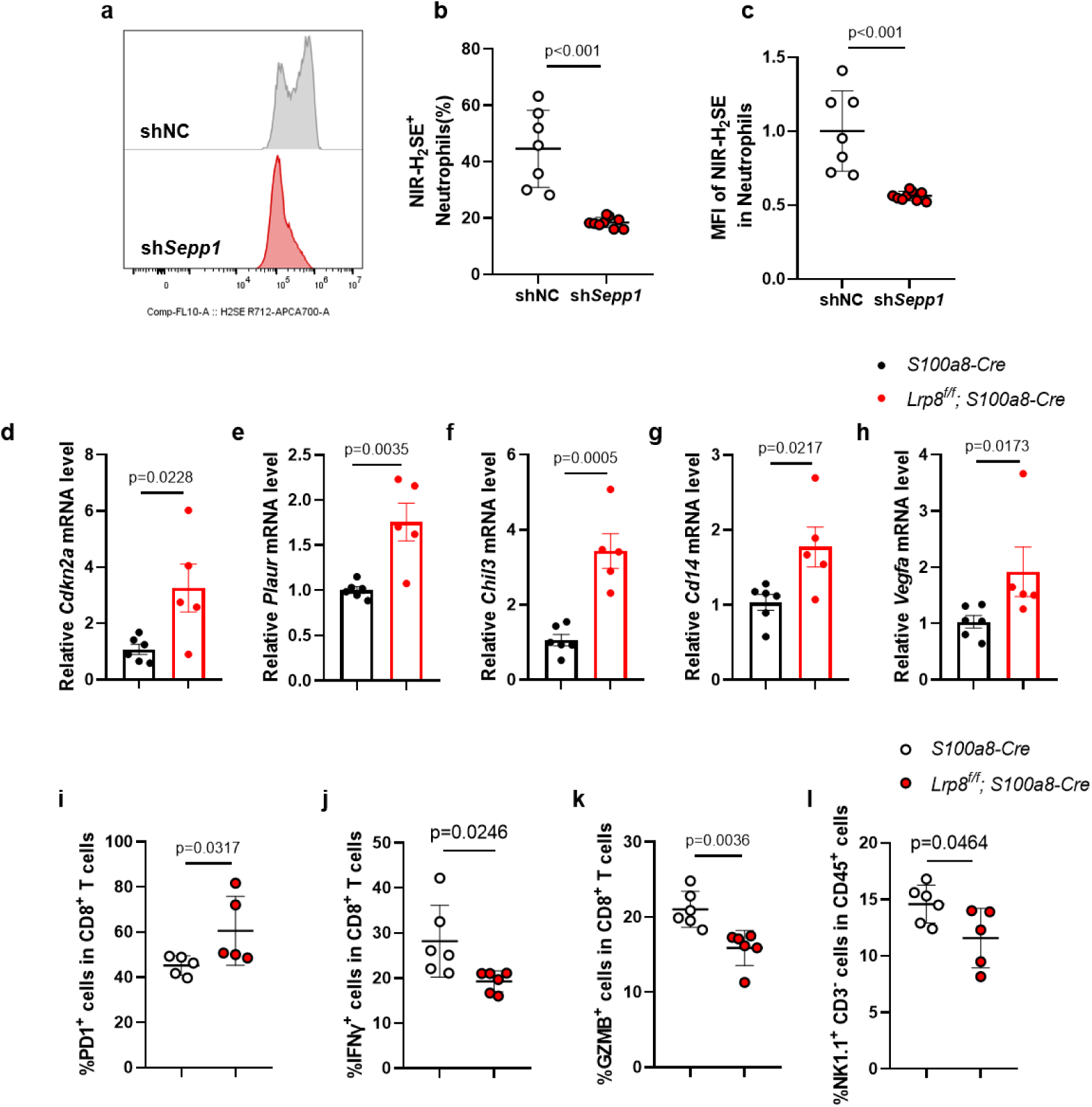
Lrp8-mediated uptake of Sepp1 by neutrophils potentiates intracellular hydrogen selenide levels (related to Figure 6). (a – c) Flow cytometric analysis of neutrophils isolated from shNC and sh*Sepp1* tumors. (a) Representative histogram showing fluorescence by NIR-H_2_SE, a hydrogen selenide probe, in neutrophils. Quantification of the percentage of NIR-H_2_SE^+^ neutrophils (b) and mean fluorescence intensity (MFI) of NIR-H_2_SE in neutrophils (c), showing significantly decreased hydrogen selenide levels in neutrophils from sh*Sepp1* tumors compared to those from shNC controls (n = 7–9 per group). (d, e)Quantitative RT-PCR analysis of senescence-associated markers *Cdkn2a* and *Plaur* in tumor-infiltrating neutrophils (n = 5–6 per group). (f-h) Analysis of *Chil3*, *Cd14*, and *Vegfa* mRNA levels in TINs from Lrp8^ΔNeu^ and control mice (n = 5–6 per group). (i–k) Functional analysis of CD8^+^ T cells in tumors, demonstrating increased PD-1 expression (i), decreased interferon gamma (IFNγ) levels (j), and decreased granzyme B (GZMB) levels (k) in Lrp8^ΔNeu^ mice compared to controls (n = 5–6 per group). (l) The percentage of natural killer (NK) cells (NK1.1^+^ CD3^−^) was significantly decreased in the tumor microenvironment of Lrp8^ΔNeu^ mice compared control animals (n = 6 per group). Data are presented as mean ± SEM; statistical significance was determined using two-tailed unpaired Student’s t-tests.

**Supplementary Figure 11:**
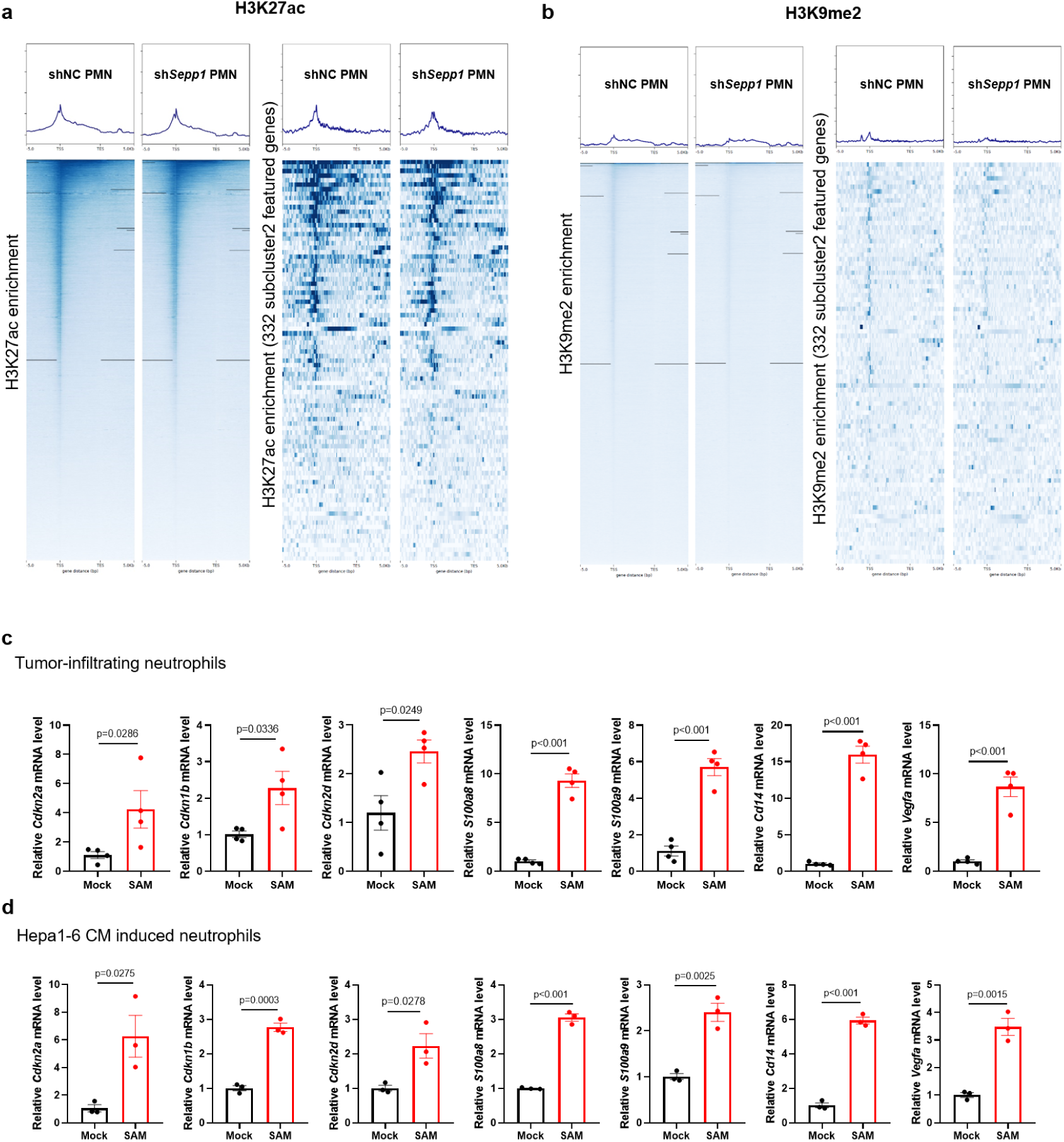
Sepp1 regulates neutrophil H3K4me3 and senescence via hydrogen selenide (related to Figure 7). (a) Heatmaps and enrichment profiles of acetylation histone H3 on lysine 27 (H3K27ac) in neutrophils isolated from tumors in shNC and sh*Sepp1* mice. The left panel shows genome-wide H3K27ac enrichment profiles in shNC and sh*Sepp1* neutrophils. The right panel highlights H3K27ac enrichment in the featured genes within the senescent-like Subcluster 2. (b) Heatmaps and enrichment profiles of demethylation of histone H3 on lysine 9 (H3K9me2) in neutrophils isolated from tumors in shNC and sh*Sepp1* mice. The left panel shows genome-wide H3K9me2 enrichment profiles in shNC and sh*Sepp1* neutrophils. The right panel highlights H3K9me2 enrichment in the featured genes within the senescent-like Subcluster 2. (c) Adding SAM to the culture medium notably upregulated the expression of *Cdkn2a*, *Cdkn1b*, *Cdkn2d*, *S100a8/9*, *Cd14* and *Vegfa* in tumor-infiltrating neutrophils. (d) Adding SAM to the culture medium notably upregulated the expression of *Cdkn2a*, *Cdkn1b*, *Cdkn2d*, *S100a8/9*, *Cd14* and *Vegfa* in Hepa1-6 CM induced neutrophils. Data are presented as mean ± SEM, with statistical significance determined by two-tailed unpaired Student’s t-tests or Mann-Whitney U-tests.

## Notes

### Competing Interest Statement

The authors have declared no competing interest.

